# MOV10 facilitates messenger RNA decay in an *N6*-methyladenosine (m^6^A) dependent manner to maintain the mouse embryonic stem cells state

**DOI:** 10.1101/2021.08.11.456030

**Authors:** Majid Mehravar, Yogesh Kumar, Moshe Olshansky, Pushkar Dakle, Michael Bullen, Varun K. Pandey, Dhiru Bansal, Craig Dent, Dhaval Hathiwala, Zhongming Zhang, Hetvi Gandhi, Alex Fulcher, Cheng Huang, Jonathan Price, Senthil Arumugam, Stephanie Ceman, Sureshkumar Balasubramanian, Brian N Papas, Marcos Morgan, Eric A Miska, Ralf B Schittenhelm, Pratibha Tripathi, Partha Pratim Das

## Abstract

*N6*-methyladenosine (m^6^A) is the most predominant internal mRNA modification in eukaryotes, recognised by its reader proteins (so-called m^6^A-readers) for regulating subsequent mRNA fates — splicing, export, localisation, decay, stability, and translation — to control several biological processes. Although a few m^6^A-readers have been identified, yet the list is incomplete. Here, we identify a new m^6^A-reader protein, Moloney leukaemia virus 10 homologue (MOV10), in the m^6^A pathway. MOV10 recognises m^6^A-containing mRNAs with a conserved GGm^6^ACU motif. Mechanistic studies uncover that MOV10 facilitates mRNA decay of its bound m^6^A-containing mRNAs in an m^6^A-dependent manner within the cytoplasmic processing bodies (P-bodies). Furthermore, MOV10 decays the Gsk-3ß mRNA through m^6^A that stabilises the ß-CATENIN expression of a WNT/ß-CATENIN signalling pathway to regulate downstream NANOG expression for maintaining the mouse embryonic stem cells (mESCs) state. Thus, our findings reveal how a newly identified m^6^A-reader, MOV10 mediates mRNA decay via m^6^A that impact embryonic stem cell biology.

## Introduction

*N6*-methyladenosine (m^6^A) is one of the most abundant and extensively studied internal reversible RNA modifications discovered in mRNAs and non-coding RNAs (lncRNAs) of eukaryotes, which plays crucial roles in splicing, export, decay/stability and translation that ultimately control various biological processes^1–4^. The m^6^A RNA methylation occurs at the *N6* position of the aromatic ring of A (adenosine), and it accounts for ∼0.2–0.6% of all A sites in mammalian mRNAs^5–8^. Epitranscriptome profiling of m^6^A using high-throughput sequencing revealed its presence mostly near the stop codon and 3’UTR of mRNAs with a ‘consensus sequence motif’ of RR(m^6^A)CH (where R refers to A/G; H refers to A/C/U)^5, 6, 9^. The deposition of m^6^A is conveyed by ‘m^6^A-methyltransferases’— METTL3 and METTL14 (m^6^A-writers) and their associated sub-units — WTAP, KIAA1429 and ZC3H13 — that form a functional m^6^A methyltransferase complex^1, 10^. Conversely, m^6^A deposition can be removed by ‘m^6^A-demethylases’ — FTO and ALKBH5 (m^6^A-erasers)^1, 10^. In addition, a group of m^6^A-binding proteins, so-called ‘m^6^A-readers’— such as YTH domain-containing proteins (YTHDF1-3) and YTHDC1,2; heterogeneous nuclear ribonucleoproteins (HNRNPs); insulin-like growth factor 2 mRNA-binding proteins (IGF2BP1-3) — have been identified that recognise m^6^A in mRNAs and govern the fate of their targeted mRNAs^11, 12^. Particularly, YTHDF family proteins (YTHDF1, YTHDF2 and YTHDF3) have been reported to play diverse molecular functions: for instance, YTHDF1 enhances mRNA translation, YTHDF2 promotes mRNA degradation, and YTHDF3 increases translation^13–15^. However, recent studies demonstrated that all YTHDFs (1-3) act combinedly and have redundant functions for binding to m^6^A-containing mRNAs and mediating their degradation^16, 17^. While other YTH domain-containing m^6^A-readers, such as YTHDFC1 perform various nuclear functions including splicing, X-chromosome inactivation and heterochromatin formation^18–22^, YTHDC2 enhances translation^23^. Moreover, remaining m^6^A-readers—HNRNPs (HNRNPA2B1, HNRNPC) facilitate pre-mRNA processing through m^6^A-switch^24, 25^; IGF2BPs (1-3) recognise m^6^A-containing mRNAs and promote their stability and translation^26^. Nonetheless, current knowledge regarding molecular and biological functions of m^6^A-readers is incomplete due to identifying a limited number of m^6^A-readers. Therefore, we have expanded the search to identify new m^6^A-reader(s) in mouse embryonic stem cells (mESCs) and how they regulate mRNA fate to maintain the mESC state.

Here, we present Moloney leukaemia virus 10 homologue (MOV10) as a novel m^6^A-reader protein. MOV10 is an RNA helicase^27^, which localises mainly at cytoplasmic processing bodies (P-bodies) for mRNA degradation and translational repression^28–30^. Nevertheless, the roles of MOV10 in m^6^A-mediated mRNA regulation have not been reported previously. This study shows that MOV10 recognises m^6^A-containing mRNAs with a conserved GGm^6^ACU motif and promotes their decay/destabilisation within the P-bodies in an m^6^A-dependent manner. Moreover, MOV10 precisely facilitates Gsk-3ß mRNA decay via m^6^A, which stabilises the ß-CATENIN expression (of a WNT/ß-CATENIN pathway) to retain downstream expression of NANOG for the mESC state maintenance. Hence, our findings provide mechanistic insights into how a new m^6^A-reader protein, MOV10, regulates mRNA decay in an m^6^A dependent manner and maintains the ESC state.

## Results

### MOV10 identified as an m^6^A-binding/reader protein that selectively binds to m^6^A-containing mRNAs

To identify potential m^6^A-reader proteins, we conducted m^6^A RNA pull-down with mESCs lysate followed by quantitative mass spectrometry-based proteomics (Fig. 1a, Extended Data Fig. 1a). First, we designed biotinylated single-stranded (ss) RNA probes— containing either three repeats of an m^6^A consensus motif of GGm^6^ACU (with methylated adenosine [A]) or three repeats of GGACU (with unmethylated adenosine [A], used as a control) (Extended Data Fig. 1a). Next, these biotinylated RNA probes were incubated with mESCs lysate, and streptavidin beads were used to pull down the RNA probes-bound proteins, which were subsequently identified through quantitative mass spectrometry-based proteomics (Fig. 1a, Extended Data Fig. 1a). The potential m^6^A-reader proteins were identified based on their enriched binding to the methylated GGm^6^ACU RNA probe over the unmethylated GGACU RNA probe (Log2FC (GGm^6^ACU/GGACU) >1; *p*-values < 0.05) (Fig. 1, Extended Data Fig. 1a, Supplementary Table 1). MOV10 was identified as one of the potential m^6^A-readers (Fig. 1a). Besides, other well-known m^6^A-readers, such as YTHDF1-3 and IGF2BP1-3, were identified in mESCs (Fig. 1a), indicating the m^6^A RNA pull-down approach was successful. Further, we confirmed that MOV10 and known m^6^A-readers (YTHDF2 and IGF2BP1,2) are preferentially bound to the methylated GGm^6^ACU RNA probe compared to the unmethylated GGACU RNA probe (Fig. 1b).

**Fig. 1.**
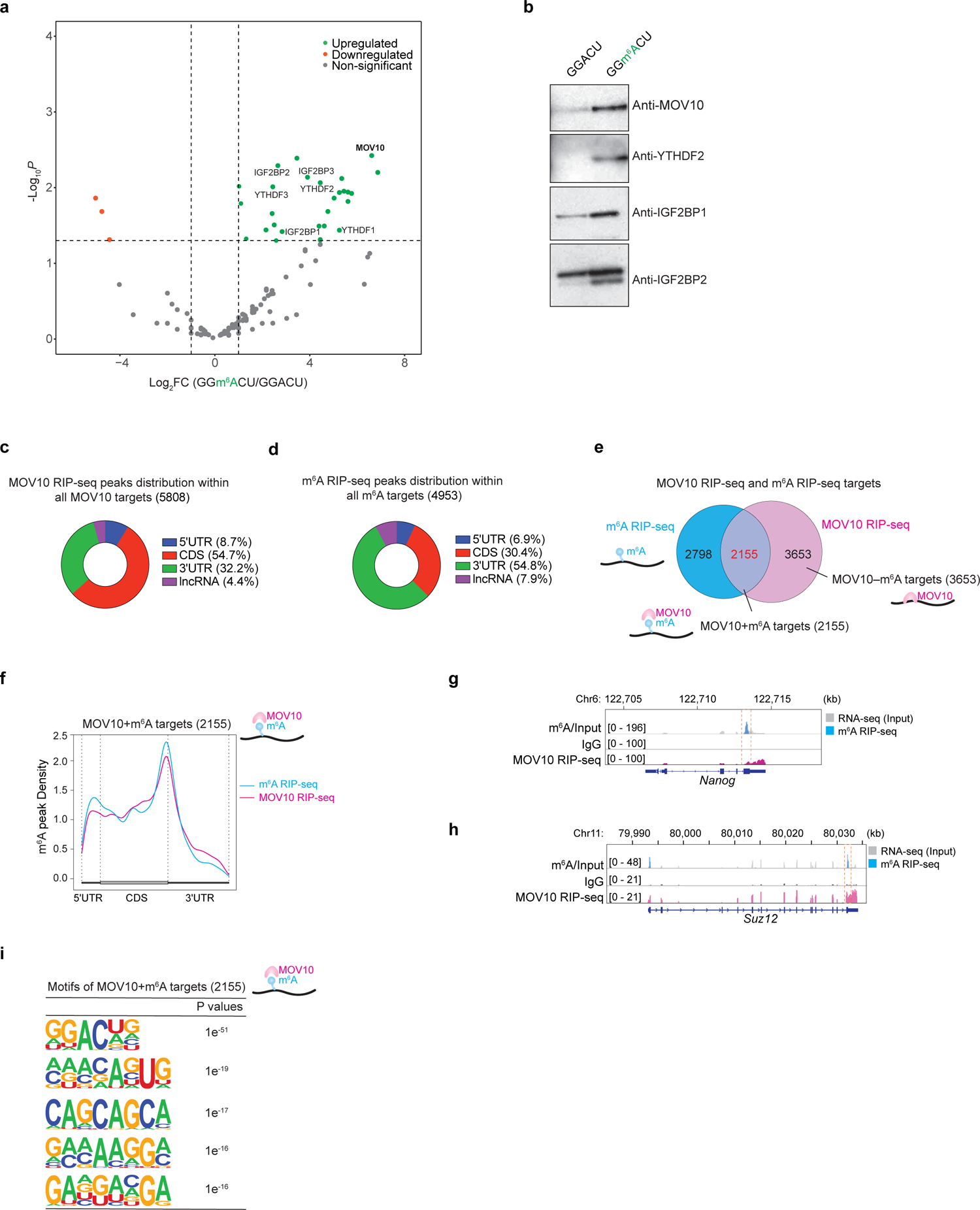
MOV10 identified as an m^6^A-binding/reader protein that selectively binds to m^6^A-containing mRNAs. a) A volcano plot represents significant m^6^A-interacting proteins that are either enriched (in green) or depleted (in red) in binding to the methylated GGm^6^ACU RNA probe over the unmethylated GGACU RNA probe (Log2FC (GGm^6^ACU/GGACU) > 1; *p*-values < 0.05). Non-significant proteins are labelled in grey. b) Western blot showing MOV10, YTHDF2, IGF2BP1 and IGF2BP2 preferentially bind to the methylated GGm^6^ACU RNA probe compared to the unmethylated GGACU RNA probe. c) Pie-chart distribution of MOV10 binding sites at 5’UTRs, CDS, 3’UTRs and long non-coding RNAs (lncRNAs) within all MOV10 targets (5,808), obtained from MOV10 RIP-seq. d) Pie-chart distribution of m^6^A sites at 5’UTRs, CDS, 3’UTRs and lncRNAs within all m^6^A-containing targets (4,953), acquired through m^6^A RIP-seq. e) Venn diagram display common and unique binding RNA targets of MOV10 and m^6^A — i.e., MOV10-bound targets with m^6^A (MOV10+m^6^A targets—2,155), MOV10-bound targets without m^6^A (MOV10-m^6^A targets—3,655) and m^6^A-containing only targets (2,798). f) Metagene plot illustrating the overlap of distribution of MOV10 RIP-seq peaks and m^6^A RIP-seq peaks in MOV10+m^6^A targets. g, h) m^6^A RIP-seq tracks showing enrichment of m^6^A over A (from RNA-seq, used as input/control) in mRNA targets; MOV10 RIP-seq tracks displaying enriched binding of MOV10 over IgG (used as a control) in mRNA targets. The dotted box highlights overlapping m^6^A and MOV10 binding sites to mRNA targets. i) Top five motifs are shown of MOV10+m^6^A targets (2,155).

To determine MOV10-bound mRNAs, we conducted RNA immunoprecipitation (RIP) of MOV10 followed by high-throughput sequencing (RIP-seq), which identified 5,808 MOV10-bound mRNA targets in mESCs (Fig. 1c, Extended Data Fig. 1b). We also performed m^6^A-seq from mESCs that identified 4,953 m^6^A-containing mRNA targets (Fig. 1d, Extended Data Fig. 1c), out of which 2,155 (43.5%) m^6^A-mRNAs were MOV10 targets (Fig. 1e). Thus, 2,155 mRNAs were considered as MOV10-bound m^6^A-containing targets, termed ‘MOV10+m^6^A targets’ (MOV10 with m^6^A targets) (Fig. 1e). However, a set of MOV10-bound mRNA targets (3,653 out of 5,808) did not contain any m^6^A and were named ‘MOV10-m^6^A targets’ (MOV10 without m^6^A targets) (Fig. 1e). A similar result has been reported for YTHDF2 m^6^A-reader, where YTHDF2 binds around half of its mRNA targets containing m^6^A ^13^. We also identified ‘MOV10 non-targets’ (7,764), where MOV10 does not bind to any of these mRNAs that are expressed in mESCs. Further analyses demonstrated that a majority of the MOV10 binding sites are located at the 3’ untranslated regions (3’UTRs) and the coding regions (CDS) of mRNAs, which coincide with overall m^6^A distribution (Figs. 1c, d). Particularly, MOV10+m^6^A targets (2,155) exhibited overlapping distribution of MOV10 binding sites and m^6^A sites at the 3’UTRs and CDS (Figs. 1f-h, Extended Data Fig. 1d). Moreover, motif analysis showed that only MOV10+m^6^A targets (2,155) consist of a GGm^6^ACU consensus motif compared to MOV10 non-targets (7,764) (Fig. 1i, Extended Data Fig. 1e). Nonetheless, MOV10-m6A targets (3,653) were excluded for motif analysis as this set of targets does not comprise any m^6^A sites. Altogether, these data suggest that MOV10 is an m^6^A-binding/reader protein, and it selectively binds to m^6^A-containing mRNAs.

### MOV10 destabilises its mRNA targets in an m^6^A-dependent manner

To dissect the functions of MOV10 in mESCs, we created *Mov10 knockout (KO)* mESC line (Extended Data Fig. 2a-c). m^6^A-seq (which assess m^6^A profiling transcriptome-wide)^9^ and RNA-seq (which measure mRNA expressions transcriptome-wide) data were generated from wild-type and *Mov10 KO* mESC lines. m^6^A-seq displayed a marked reduction of m^6^A levels/intensities of MOV10+m^6^A targets in *Mov10 KO* compared to wild-type (Fig. 2a, c). In contrast, MOV10 non-targets did not significantly alter m^6^A levels between *Mov10 KO* and wild-type mESCs (Fig. 2b, d). Of note, we excluded MOV10-m^6^A targets to measure m^6^A changes in *Mov10 KO,* since this set of MOV10 mRNA targets is devoid of any m^6^A sites (Fig. 1e). Nonetheless, we did not observe substantial changes of m^6^A levels of ‘all’ m^6^A-containing targets in *Mov10 KO* compared to wild-type mESCs (Extended Data Fig. 2d), indicating that MOV10 modulates m^6^A levels specifically to its bound m^6^A-containing targets. Further, differential gene expression analysis (RNA-seq) demonstrated upregulation of only MOV10+m^6^A targets in *Mov10 KO* compared to wild-type. Whereas no significant changes were observed of MOV10-m^6^A targets and MOV10 non-targets (Fig. 2e and Extended Data Fig. 2e).

**Fig. 2.**
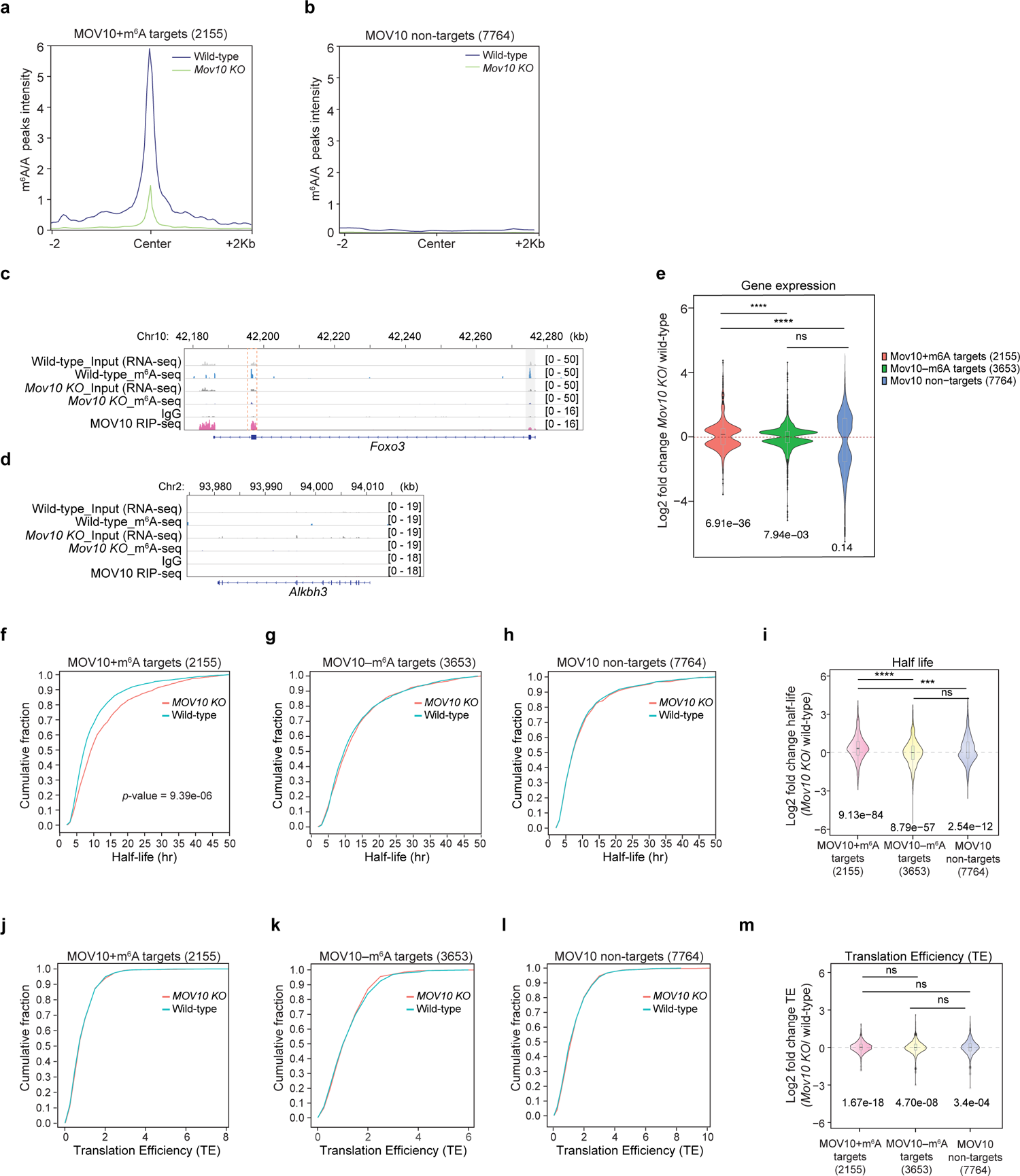
MOV10 destabilises its mRNA targets in an m^6^A-dependent manner. a, b) m^6^A/A peak intensities of MOV10+m^6^A targets (2,155) and MOV10 non-targets (7,764) in wild-type and *Mov10 KO*. c) m^6^A RIP-seq and RNA-seq (control) tracks showing the enrichment of m^6^A in mRNA target. RIP-seq of MOV10 and IgG (control) tracks displaying the enrichment of MOV10 binding to mRNA target. Dotted box feature overlapping m^6^A and MOV10 enriched sites in a MOV10+m^6^A target, where m^6^A level is reduced in *Mov10 KO* compared to wild-type. d) A typical MOV10 non-target region show absence of MOV10 and m^6^A enriched peaks. e) A violin plot represents gene expression changes of MOV10+m^6^A targets, MOV10-m^6^A targets and MOV10 non-targets in *Mov10 KO* compared to wild-type. *P*-values of each set of targets are shown based on the Wilcoxon t-test (at the bottom). Significant differences between two sets of targets are represented through *p*-values using ANOVA (at the top); *p*\*\*\*\*< 0.0001; ns = non-significant; n = 2 replicates. f-h) Cumulative plots exhibiting HLs of MOV10+m^6^A targets (2,155) (f), MOV10-m^6^A targets (3,653) (g), and MOV10 non-targets (7,764) (h) in wild-type v. *Mov10 KO*. Only significant *p*-values that represent the HL changes in *Mov10 KO* v. wild-type are shown in each graph; n = 2 replicates. i) A violin plot presenting log2 fold change of HLs of MOV10+m^6^A targets, MOV10-m^6^A targets and MOV10 non-targets in *Mov10 KO* compared to wild-type. *P*-values are shown of each set of targets (Wilcoxon t-test) (at the bottom). Significant changes among two sets of targets are represented through *p*-values (ANOVA test) (at the top); *p*\*\*\*\*< 0.0001, *p*\*\*\*< 0.001, ns = non-significant; n = 2 replicates. j-l) Cumulative plots displaying TE of MOV10+m^6^A targets (j), MOV10-m^6^A targets (k), MOV10 non-targets (l) in wild-type v. *Mov10 KO*. m) A violin plot viewing log2 fold change of TE of MOV10+m^6^A targets, MOV10-m^6^A and MOV10 non-targets in *Mov10 KO* compared to wild-type. *P*-values are shown of each set of targets using the Wilcoxon t-test (at the bottom). Changes among two sets of targets are ns = non-significant (ANOVA test); n = 3 replicates.

To determine whether MOV10 regulates its mRNA targets post-transcriptionally, we measured half-life (HL) (that evaluates the mRNA decay/ stability)^13^ and translation efficiency (TE) (that quantifies the rate of mRNA translation)^31^ of MOV10+m^6^A targets, MOV10-m^6^A targets, and MOV10 non-targets in *Mov10 KO* and wild-type. We found that HL was significantly increased of MOV10+m^6^A targets in the absence of *Mov10* (Figs. 2f, i), but half-lives (HLs) of MOV10-m^6^A targets and MOV10 non-targets were unchanged in *Mov10 KO* compared to wild-type mESCs (Fig. 2g, h, i). Further, TE of MOV10+m^6^A targets, MOV10-m^6^A targets and MOV10 non-targets were remained the same between *Mov10 KO* and wild-type (Fig. 2j-m and Extended Data Fig. 2f-i). Altogether, these findings implying that MOV10 destabilises/decays its mRNA targets in an m^6^A-dependent manner.

### MOV10 interactome establishes diverse protein networks yet maintain the P-bodies integrity

To dissect the potential molecular functions of MOV10 by which it regulates its m^6^A-containing targets for decay, we conducted MOV10 Immunoprecipitation (IP) followed by quantitative mass spectrometry (IP-MS) to identify MOV10-interacting proteins in mESCs (Extended Data Fig. 3a). Several high-confidence MOV10-interacting proteins were identified (Log2FC > 1; *p*-values < 0.05) (Supplemental Table 2). Gene Ontology (GO) analysis demonstrated that MOV10-interacting proteins are conceivably involved in various biological functions, such as mRNA processing, mRNA splicing, mRNA polyadenylation, mRNA transport, mRNA stability and translation (Fig. 3a). Also, they are predominantly enriched in ribonucleoprotein (RNP) complex, mRNA P-bodies, nucleus, cleavage and polyadenylation specificity factors (CPSFs) complex and spliceosome complex (Fig. 3b). Indeed, we showed that MOV10 has nuclear and cytoplasmic localisation, including P-bodies (Fig. 3d). Notably, we found that several P-body proteins (IGF2BP1, IGF2BP2, IGF2BP3, EDC4, ZCCHC3, YTHDF2, YTHDC1, ELAVL1, ELAVL2, AGO2, EIF4A3, DHX9, DHX15, DHX21, DHX30, APOBEC3F, TUT4 and UPF1) as well as splicing factors (SRSFs, U2AF2, SF3B1,3), HNRNPs, CPSFs and RNA-binding proteins (PABPC1, FXR1, 2 etc.) are the major MOV10-interacting partners in mESCs (Fig. 3c; Extended Data Fig. 3b-f).

**Fig. 3.**
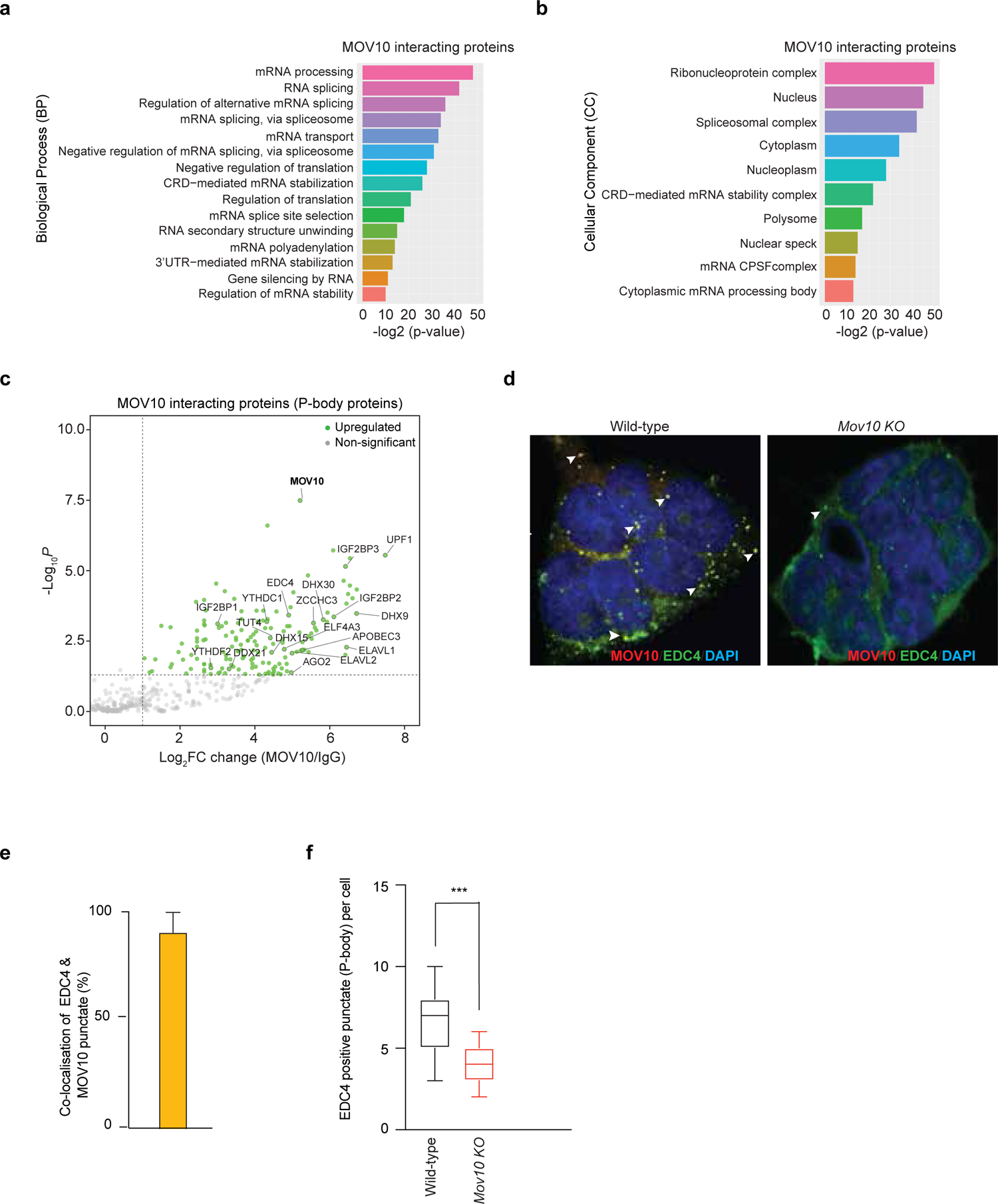
MOV10 interactome establishes diverse protein networks yet maintain the P-bodies integrity. a, b) Gene Ontology (GO) term analysis (biological process) (a) and (cellular components) (b) of significantly enriched MOV10-interacting partners that are obtain through MOV10 Immunoprecipitation followed by quantitative mass spectrometry (IP-MS). c) Significantly enriched MOV10-interacting P-body proteins are shown (in green) that are identified through MOV10 IP-MS (Log2FC (MOV10/IgG) > 1; *p*-values < 0.05). Non-significant proteins are labelled in grey, n *=* 2 replicates. d) Immunostaining showing sub-cellular co-localisation of MOV10 and EDC4 (as a P-body marker) as punctate structures. Loss of *Mov10* shows a significant reduction of EDC4+ve punctate structures. e) Quantification of sub-cellular co-localisation of MOV10 and EDC4. f) EDC4+ve punctuates in *Mov10 KO* v. wild-type. *P*-values were calculated using ANOVA; *p*\*\*\*< 0.001; n = 3 replicates, including 90 cells.

Because CPSFs, PABPC1, TUT4 were identified as MOV10-interacting partners (Fig. 3a, Extended Data Fig. 3c, e), they are critical for mRNA 3’-end processing and polyadenylation signals/length^32–34;^ we measured poly(A) length changes in the absence of *Mov10*. No significant alterations of poly(A) length of MOV10+m^6^A targets, MOV10-m^6^A targets and MOV10 non-targets were observed in *Mov10 KO* compared to wild-type (Extended Data Fig. 3g-i). Since CPSFs, HNRNPs and splicing proteins (SRSFs, U2AF2, SF3B1,3) were also identified as MOV10-interacting proteins (Extended Data Fig. 3b-d), and they are vital for splicing^32, 35^, we examined MOV10 functions in splicing. Although we observed a significant correlation between individual m^6^A sites and individual splice-sites strength estimates (SSE)/usage^36^, no alternations of SSE/usage were observed around the m^6^A sites in *Mov10 KO* compared to wild-type (Extended Data Fig. 3j). Further, we found that MOV10 is associated with EIF4A3 (a translation initiation factor) but doesn’t affect the TE of its m^6^A-containing targets (Fig. 2j-m; Fig. 3c). Nonetheless, we demonstrated that MOV10 precisely decays its m^6^A-containing mRNA targets (Fig. 2f-i). This perhaps happens through nonsense-mediated mRNA decay (NMD), as MOV10 strongly interacts with UPF1 (Fig. 3a, Supplemental Table 2), and a previous study showed that MOV10 mediates UPF1-directed NMD^29^.

MOV10 was identified mainly as a P-body protein^37, 38^, and the majority of its interacting partners were P-body proteins (Fig. 3c, Supplemental Table 2). Thus, we asked whether MOV10 maintains the P-bodies integrity. P-bodies are cytoplasmic RNP granules comprising mRNAs and proteins that act as a site for regulating the fate of mRNAs (mRNA decay, mRNA surveillance and translational repression)^30^. We assessed the cellular localisation of MOV10 by immunostaining along with P-bodies marker EDC4^32, 33^. It revealed significant co-localisation (93%) of MOV10 and EDC4 as typical punctate structures of P-bodies (Fig. 3d, e). This was consistent with the proteomic analysis, which exhibited a physical association between MOV10 and EDC4 (Fig. 3c). However, loss of *Mov10* showed a marked reduction of EDC4+ve punctate structures of P-bodies (Fig. 3d, f), suggesting that MOV10 preserve the P-bodies integrity in mESCs.

### MOV10 destabilises its m^6^A-containing mRNA targets mainly within the P-bodies

We demonstrated MOV10 destabilises its m^6^A-containing mRNA targets (Fig. 2f, i), and it predominantly presents within P-bodies to maintain the P-bodies integrity (Fig. 3d-f). Hence, we hypothesised that MOV10 might promote the destabilisation of its m^6^A-mRNA targets within the P-bodies. To test this postulation, we first generated RIP-seq data of DDX6 (a vital component of P-bodies^30, 37^) from mESCs, which identified P-bodies associated DDX6-bound mRNA targets (Extended Data Fig. 4a, b). Of note, we found significant co-localisation (89%) between DDX6 and MOV10 (Extended Data Fig. 4c). Next, we compared DDX6-bound (6,910) and MOV10-bound (5,808) mRNA targets; it revealed that 69% (4,009 out of 5,808 targets) of MOV10 mRNA targets overlapped with DDX6 mRNA targets, named ‘MOV10+DDX6 targets’ (Extended Data Fig. 4d). Moreover, the majority of the MOV10+m^6^A targets (∼89%; 1,914 out of 2,155 targets) shared with DDX6 targets (named MOV10+m^6^A+DDX6 targets); whereas ∼57% (2,095 out of 3,653 targets) of the MOV10-m^6^A targets overlapped with DDX6 targets (named MOV10-m^6^A+DDX6 targets) (Fig. 4a). Half-life (HL) calculation revealed a significant increase of HLs of MOV10+DDX6 (4,009) and MOV10+m^6^A+DDX6 (1,914) targets in *Mov10 KO* compared to wild-type (Fig. 4b, d; Extended Data Fig. 4e, g). In contrast, MOV10 only targets (1,799) (that does not overlap with DDX6 targets) showed a modest decrease of HLs in *Mov10 KO* (Extended Data Fig. 4f, g). However, MOV10-m^6^A+DDX6 targets (2,095) and MOV10 non-targets (7,764) displayed no significant alterations of their HLs between *Mov10 KO* and wild-type (Fig. 2h, Fig. 4c, d) – denoting that MOV10 destabilises its bound mRNA targets, including m^6^A-mRNA targets that are shared with DDX6-bound mRNA targets. However, TE of any of the mRNA target sets were unaltered in the absence of *Mov10* (Fig. 4e-g; Extended Data Fig. 4h-j). Taken together, these data suggest that MOV10 mediates the destabilisation/decay of its m^6^A-containing mRNA targets within the P-bodies.

**Fig. 4.**
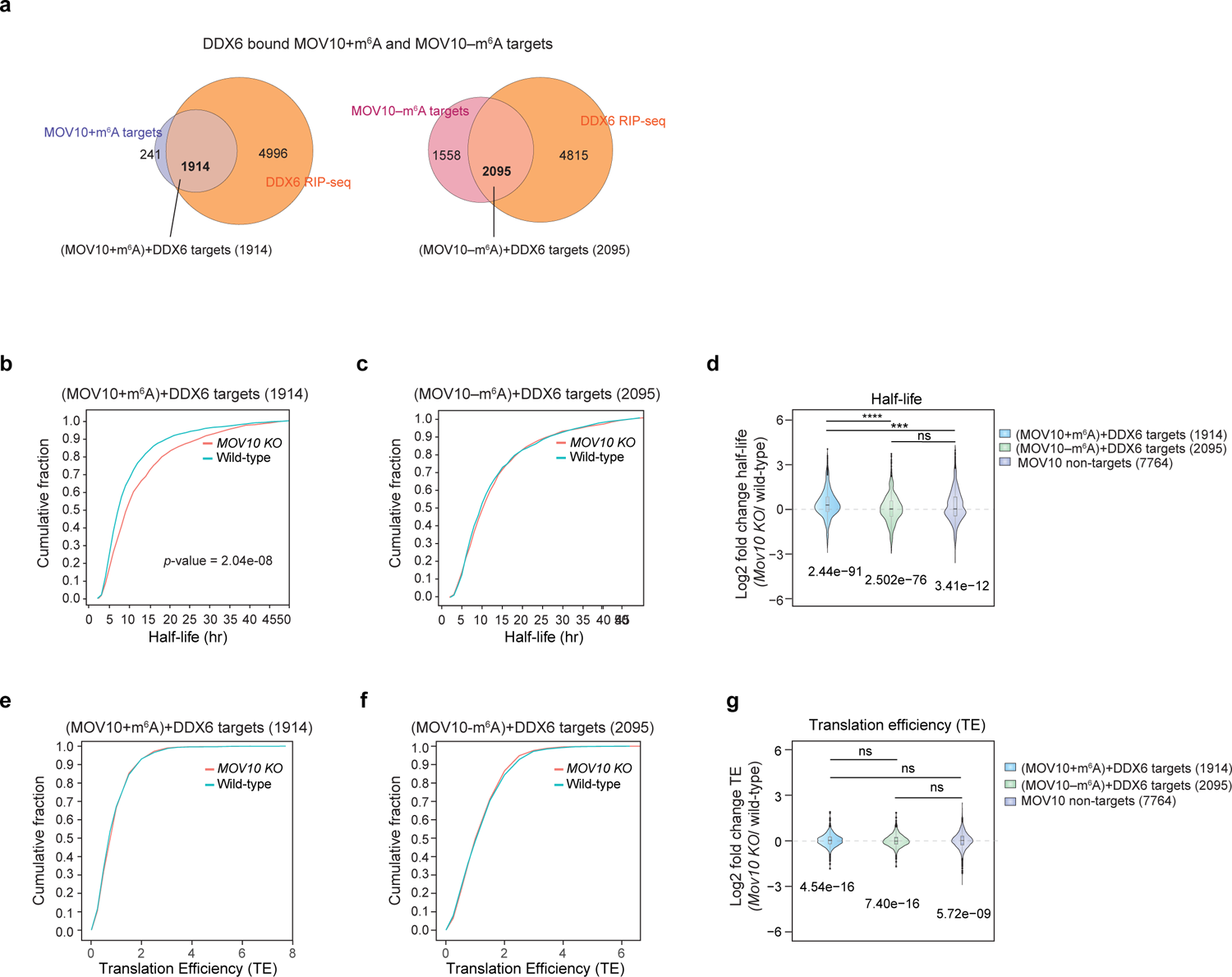
MOV10 destabilises its m^6^A-containing mRNA targets mainly within the P-bodies. a) Venn diagrams represent overlap targets between—MOV10+m^6^A targets (2,155) and DDX6 targets (6,910) (left panel), as well as MOV10-m6A targets (3,653) and DDX6 targets (6,910) (right panel). This exhibits MOV10+m^6^A+DDX6 targets (1,914) and MOV10-m^6^A+DDX6 targets (2,095). b, c) Cumulative plots showing HLs of MOV10+m^6^A+DDX6 targets (1,914) and MOV10-m^6^A+DDX6 targets (2,095). Only significant *p*-values that represent the HL changes in *Mov10 KO* v. wild-type are shown in each graph; n = 2 replicates. d) A violin plot exhibiting log2 fold change of HLs of MOV10+m^6^A+DDX6 targets, MOV10-m^6^A+DDX6 and MOV10 non-targets in *Mov10 KO* compared to wild-type. *P*-values are shown for each set of targets using the Wilcoxon t-test (at the bottom). Significant changes among two sets of targets are displayed through *p*-values (ANOVA test) (at the top); *p*\*\*\*\*< 0.0001, *p*\*\*\*< 0.001, ns = non-significant; n = 2 replicates. e, f) Cumulative plots demonstrating TE of MOV10+m^6^A+DDX6 targets (1,914) (e) and MOV10-m^6^A+DDX6 targets (2,095) (f) in wild-type v. *Mov10 KO*. g) Violin plot showing log2 fold change of TE of MOV10+m^6^A+DDX6 targets, MOV10-m^6^A+DDX6 and MOV10 non-targets in *Mov10 KO* compared to wild-type. *P*-values are indicated of each set of targets (at the bottom). Alterations among two sets of targets are ns= non-significant (ANOVA test); n=3 replicates.

### MOV10 destabilises Gsk-3ß mRNA in an m^6^A-dependent manner to control the WNT/ß-CATENIN-NANOG regulatory pathway for the mESC state maintenance

mESCs typically grow as round/dome-shaped three-dimensional colonies^40^. However, *Mov10 KO* mESCs grew as monolayer and exhibited flattened cellular morphology with reduced cell-to-cell contacts (Fig. 5a), indicating that *Mov10 KO* mESCs were differentiated (i.e., losing their stemness/ESC state). To investigate the underlying mechanisms of this phenotype, we inspected HL changes of MOV10+m^6^A mRNA targets (2,155) in *Mov10 KO* compared to wild-type. We found that the HL of glycogen synthase kinase-3ß (Gsk-3ß) mRNA was markedly increased in *Mov10 KO* (Fig. 5b, c). However, the HL of Gsk-3α (homologue of Gsk-3ß) remains unchanged in *Mov10 KO* (Fig. 5b, c). GSK-3s have been implicated for their functions to maintain the mESC state^39, 40^; particularly, GSK-3ß maintain self-renewal of the mESC state ^41^. Moreover, inhibition of GSK-3 activates the canonical WNT/ß-CATENIN pathway through stabilisation of ß-CATENIN and that subsequently activates NANOG expression (an essential ESC-TF ^42^) — suggested to be the key mechanism for the maintenance of self-renewal of mESC state ^39–41, 43, 44^. Based on these findings, we enquired whether and how GSK-3 might regulate NANOG expression in the absence of *Mov10*. To examine this, we measured Ser9 phosphorylation of GSK-3ß that is vital for controlling the ß-CATENIN degradation of the WNT/ß-CATENIN signalling pathway for mESC state maintenance ^39^. It revealed that activation of GSK-3ß Ser9 phosphorylation in the absence of *Mov10* reduced ß-CATENIN expression and following NANOG expression at the protein level (Fig. 5e-h), which ultimately causes differentiation phenotype of *Mov10 KO* mESCs (Fig. 5a). However, the total protein level of GSK-3ß remains unchanged in *Mov10 KO* compared to wild-type (Fig. 5e), suggesting that the total protein level of GSK-3ß is perhaps maintained through other pathways as well.

**Fig. 5.**
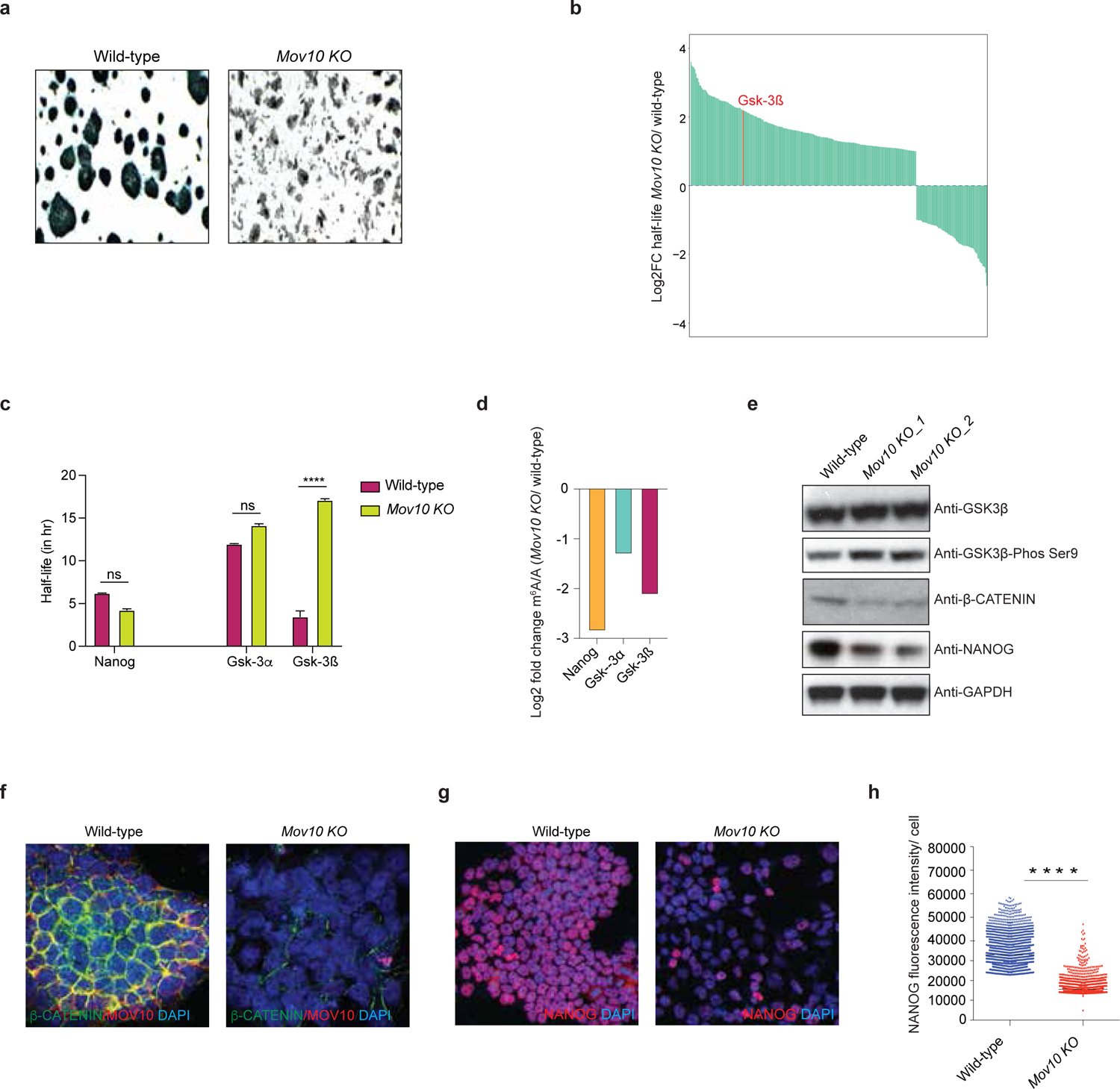
MOV10 destabilises Gsk-3ß mRNA in an m^6^A-dependent manner to control the WNT/ß-CATENIN-NANOG regulatory pathway for the mESC state maintenance. a) AP staining displays wild-type and *Mov10 KO* mESCs morphology. b) Log2 fold change of HLs of MOV10+m^6^A targets (2,155) in *Mov10 KO* compared to wild-type. c) Bar plot showing HLs of Gsk-3α, Gsk-3β and Nanog transcripts in wild-type and *Mov10 KO* mESC lines. d) Changes of m^6^A/A levels of Gsk-3α, Gsk-3ß and Nanog mRNAs in *Mov10 KO* compared to wild-type are shown. e) Protein levels of GSK-3β, GSK-3β-Phos Ser9, β-CATENIN and NANOG are displayed (Western blot) in wild-type and *Mov10 KO* mESC lines. GAPDH was used as a control. f, g) Immunostaining of MOV10, β-CATENIN and NANOG are presented in wild-type and *Mov10 KO* mESC lines. h) Quantification of NANOG protein expression (fluorescence intensity) in wild-type and *Mov10 KO* mESCs via immunostaining. *P*-values were calculated using ANOVA. \*\*\*\**p*< 0.0001; n = 3 replicates, including 100 cells.

To check whether MOV10 regulates the decay of Gsk-3ß mRNA in an m^6^A-dependent manner, we measured the m^6^A level of Gsk-3ß mRNA in the absence of *Mov10*. It showed a reduction of m^6^A level of Gsk3ß mRNA in *Mov10 KO* compared to wild-type (Fig. 5d; Extended Data Fig. 6a). Combining this with the increased HL of Gsk-3ß mRNA in the absence of *Mov10* implies that MOV10 destabilises Gsk-3ß mRNA via m^6^A. Notably, the m^6^A level was decreased of Nanog mRNA, but its HL remained the same in *Mov10 KO* compared to wild-type (Fig. 5d; Extended Data Fig. 5b), indicating that MOV10 does not regulate Nanog mRNA through m^6^A. Instead, MOV10 regulates NANOG (at the protein level) via WNT/ß-CATENIN pathway. Altogether, these findings suggest that MOV10 destabilises/decays Gsk-3ß mRNA in an m^6^A-dependent manner, stabilising the ß-CATENIN expression of a WNT/ß-CATENIN pathway for retaining downstream NANOG expression to maintain the mESC state (Fig. 6).

**Fig. 6.**
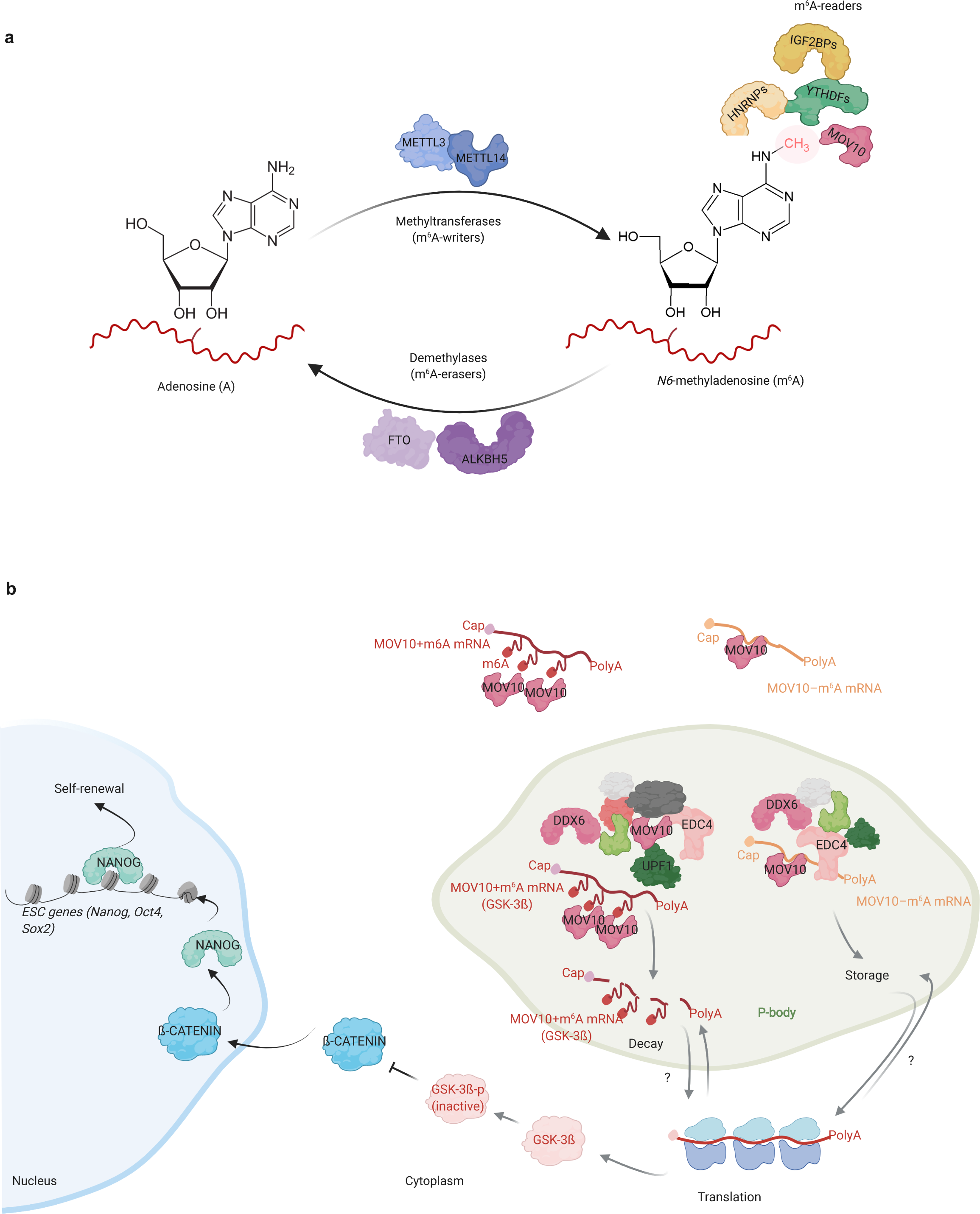
A model represents the detailed molecular functions of MOV10 to maintain the mESC state. a) The proposed model describes MOV10 as a new m^6^A-binding/reader protein in the m^6^A pathway. Other known m^6^A readers (YTHDFs, IGF2BPs, HNRNPs), and m^6^A-writers (METTL3, METTL14) & m^6^A-erasers (FTO, ALKBH5) are shown. b) We demonstrate that almost half of the total m^6^A-containing mRNA targets bind to MOV10, called MOV10+m^6^A targets; the remaining MOV10 bound mRNA targets do not contain any m^6^A, termed as MOV10-m^6^A targets. The majority of the MOV10+m^6^A targets localised within the P-bodies and is destabilised by MOV10. However, a fraction of MOV10-m^6^A targets is localised within the P-bodies and not undergoes mRNA decay via MOV10. Furthermore, our findings reveal a mechanism by which MOV10 decays Gsk-3ß mRNA (a MOV10+m^6^A target) via m^6^A that stabilises the ß-CATENIN expression of a WNT/ß-CATENIN pathway for retaining downstream NANOG expression to maintain the mESC state. (?) represent these pathways are not entirely understood yet.

## Discussion

Here, we identified MOV10 as a new m^6^A-binding/reader protein (Fig. 1) and showed that almost half of the total m^6^A-containing mRNA targets bind to MOV10 and have a consensus m^6^A motif, i.e., GGm^6^ACU (Fig. 1). Furthermore, our comprehensive molecular studies demonstrated that MOV10 facilitates mRNA decay of its bound m^6^A-containing mRNA targets, predominately within the P-bodies (Figs. 2, 3, 4).

So far, a handful of m^6^A-readers have been identified that belongs to mainly three families: YTH family members (including YTHDF1-3, YTHDC1 and YTHDC2)^12^, IGF2BP family members (including IGF2BP1-3)^26^ and HNRNP family members (including HNRNPC, HNRNPG, HNRNPA2B1)^24, 25^. These sets of m^6^A-reader proteins have diverse molecular functions, including mRNA decay/stability, translation, splicing, pre-mRNA processing, X-chromosome inactivation and heterochromatin formation^12, 18–20, 25^. Here, our study adds an m^6^A-reader protein, MOV10, into the above listing catalogue of m^6^A-readers and reveal its roles in mRNA decay in an m^6^A-dependent manner (Fig. 2). This is similar to the functions of YTHDF2, which also regulates the mRNA decay of its m^6^A-containing mRNA targets^13^. Moreover, recent studies determined the redundant function of YTHDF1-3 in the degradation/decay of their m^6^A-containing mRNA targets for mESCs differentiation^17^. Likewise, a unified function of YTHDF1-3 was also reported in the degradation of their m^6^A-containing mRNAs in cancer cells ^16^. However, both of these studies concluded that YTHDF1-3 does not affect the TE of their m^6^A-containing mRNAs^13, 16, 17^. Similarly, we also found that MOV10 does not affect the TE of its m^6^A-containing targets (Fig. 2). Though both MOV10 and YTHDFs decay their m^6^A-containing mRNA targets, the underlying mechanisms could be different^13^ (Fig. 2). For instance, YTHDF1-3 physically interacts with many CNOT components of the CCR4-NOT RNA degradation complex^16^; therefore, it is thought to degrade their m^6^A-mRNA targets through deadenylation and 3’–5’ decay^16, 45^. Whereas MOV10 physically interacts with mainly P-bodies components, including UPF1, which mediate NMD (Fig. 3) – indicate that MOV10 is perhaps decaying its m^6^A-mRNA targets through NMD as described earlier^29^. Even though MOV10 is an m^6^A-reader protein, we found that it modulates m^6^A levels of its m^6^A-mRNA targets (Fig 2a), and this perhaps happens due to the helicase activity of MOV10^29^. Likewise, a previous report demonstrated that YTHDF2 also regulates m^6^A levels of its m^6^A-mRNA targets, but the mechanism was not defined ^13, 46^. Contrariwise, IGF2BP family m^6^A-readers (IGF2BP1-3) have been shown promoting mRNA stability and translation in cancer cells; but alterations of m^6^A levels in their m^6^A-mRNA targets were not reported^26^. Altogether, it reveals that different m^6^A-reader proteins have distinct molecular functions for gene regulation at diverse cellular settings.

In conclusion, we demonstrate that MOV10 is a new m^6^A-reader protein in the m^6^A pathway, and it regulates mRNA decay of its bound m^6^A-containing mRNA targets. Furthermore, we uncover a mechanism by which MOV10 decays Gsk-3ß mRNA via m^6^A that stabilises the WNT/ß-CATENIN-NANOG regulatory pathway ^39, 41, 44^ to maintain the self-renewal of mESC state (Figs. 5, 6).

## Methods

### Mouse embryonic stem cells (mESCs) culture

mESCs were cultured in mESC media that contains — DMEM (Dulbecco’s modified Eagle’s medium) (Thermo Fisher Scientific) supplemented with 15% fetal calf serum (FCS) (Thermo Fisher Scientific), 0.1 mM β-mercaptoethanol (Sigma-Aldrich), 2 mM L-glutamine (Thermo Fisher Scientific), 0.1 mM nonessential amino acid (NEAA) (Thermo Fisher Scientific), 1% of nucleoside mix (Merck Millipore), 1000 U/mL recombinant leukemia inhibitory factor (LIF/ESGRO) (Merck Millipore) and 50 U/mL Penicillin–Streptomycin (Thermo Fisher Scientific) — at 37 ^°^C, 5% CO_2_, as described previously^47–49^.

### Antibodies

MOV10: ab176687 (Abcam) and A301-571A (Bethyl Laboratories); ß-CATENIN: 05-665-25UG (Merck); NANOG: A300-397A (Bethyl Laboratories); EDC4: ab72408 (Abcam); DDX6: NB200-191 (Novus Biologicals) and A300-461A (Bethyl Laboratories); GSK3ß-Ser9 Phos: 5558 (Cell Signaling Technology); GSK3ß: 9315 (Cell Signaling Technology); m^6^A: 202 003 (Synaptic Systems).

### Generation of *Mov10 KO* mESC lines using CRISPR/Cas9

Paired sgRNAs were designed to delete exons 1–4 of the *Mov 10* gene. These sgRNAs were cloned into the lentiguide-mCherry-Cas9 plasmid, using Golden Gate Cloning approach as mentioned previously^48^. 2 million mESCs were transfected with 1 ug of each plasmid carry sgRNAs-mCherry-Cas9, using lipofectamine 3000 (Thermo Fisher Scientific). The next day, mCherry-positive mESCs were sorted by FACS and plated on a 10 cm plate at very low density for single-cell clone picking. After 5–6 days, individual colonies (derived from single-cell) were picked, expanded and genotyped using genomic PCR to detect the ‘homozygous/biallelic deletion’ of *Mov10 KO* mESC colonies. Selected *Mov10 KO* mESC lines were chosen, grown and confirmed by Sanger sequencing and Western blot.

### Generation of MOV10-FB tagged mESC lines

The open reading frame (ORF) of mouse *Mov10* was cloned into BamHI-digested pEF1a-FlagBiotin (FB)-puromycin vector, using Gibson Assembly^®^ Master Mix (NEB). Positive clones were analysed by Sanger sequencing. 2 µg of *Mov10*-FB construct was electroporated into 5×10^6^ J1 wild-type mESCs, which constitutively express BirA ligase (neomycin). The electroporated cells were plated on a 10 cm dish with mESC media. After 24 hours, media was replaced with fresh mESC media containing 1 μg/mL of puromycin (Sigma-Aldrich) and 1 μg/mL of neomycin (Sigma-Aldrich), and cells were selected for 4–5 days. Individual mESC colonies were picked, expanded and tested by Western blot using either streptavidin-HRP antibody (GE Healthcare) (dilution 1:2000 in 5% BSA) or specific antibody against MOV10 to detect the clones that expressed MOV10-FB. MOV10-FB mESC lines that expressed comparable levels of endogenous MOV10 protein were chosen for further experiments.

### AP staining

Alkaline Phosphatase (AP) staining of mESCs was performed according to the manufacture’s protocol (Vector Black Substrate Kit, SK-5100, Vector Lab).

### Immunostaining

Cultured mESC cells (wild-type and *Mov10 KO*) were fixed with 4% paraformaldehyde and washed three times with 1X PBS. Primary antibodies were used, including MOV10 (ab176687, Abcam; A301-571A, Bethyl Laboratories), DDX6 (A300-461A, Bethyl Laboratories), EDC4 (ab72408, Abcam), NANOG (A300-397A, Bethyl Laboratories), ß-CATENIN (05-665-25UG, Merck). Secondary antibodies were obtained from Jackson ImmunoResearch. Pictures were taken with a Nikon confocal microscope; analyses were performed using Image J.

### m^6^A RNA-protein pull-down

mESCs were grown at 80% confluency on a 15 cm dish, harvested and washed twice with 1X PBS (Gibco). Cell pellet was lysed in 1 mL extraction buffer (EB) (20 mM Tris-HCl (pH 7.5), 100 mM NaCl, 5 mM MgCl_2_, 0.5% NP40 with freshly added 1:100 protease inhibitor (P8340, Sigma) and 100 U of RNase inhibitor (20 U/µL, AM2696, Thermo Fischer Scientific)) for 10 min on ice and then spun at 15,000 g for 10 min at 4 °C to remove cell debris. The supernatant was collected, 500 µg of total cell lysate (250 µL; 2 µg/µL) was incubated with 200 pmol of biotinylated ssRNA probe containing three repeats of either GGm^6^ACU or GGACU in an equal volume of 250 µL of 2X TENT buffer (20 mM Tris-HCl (pH 8), 2 mM EDTA (pH 8), 500 mM NaCl, 1% (v/v) Triton X-100) with 5 µL of protease inhibitor (P8340, Sigma) and 2.5 µL of RNase inhibitor (AM2696, Thermo Fischer Scientific) for 30 min at room temperature with gentle rotation. 50 µL of streptavidin Dynabeads™ (Cat. No. 65601, Thermo Fischer Scientific) were washed twice with 0.5 mL of 1X TENT buffer and added to this mix, incubated for an additional 30 min at room temperature. Immunoprecipitated ‘protein-RNA-Dynabeads’ were washed three times with 1 mL of 1X TENT buffer. RNA probes-bound proteins were eluted in 40 µL 2X XT buffer (Bio-Rad) and subjected to quantitative MS and/or Western blots analyses.

Biotinylated ss RNA probes (IDT) that were used:

rGrG/iN6Me-rA/rCrUrGrG/iN6Me-rA/rCrUrGrG/iN6Me-rA/rCrU/3Bio/

rGrGrArCrUrGrGrArCrUrGrGrArCrU/3Bio/

### RNA immunoprecipitation sequencing (RIP-seq)

10 million mESCs were collected, washed twice with 1X PBS and lysed in 1 mL of RNA immunoprecipitation (RIP) buffer (20 mM Tris-Cl (pH 7.4), 150 mM NaCl, 1 mM EDTA, 0.5% NP-40, 1 mM DTT with freshly added 1:1000 protease inhibitor (Sigma) and 100 U of RNase inhibitor (20 U/µL, Thermo Fisher Scientific)) for 30 min on ice. The cell lysate was cleared by centrifugation at 15,000 g for 10 min at 4 °C. Subsequently, the supernatant was collected and divided into two equal volumes (0.5 mL each). 5 µg of an antibody of interest (MOV10 or DDX6) and 5 µg of anti-IgG antibody (as a control) were added to the supernatant (in 0.5 mL) and incubated for two hours at 4 °C with gentle rotation. 50 μL of Dynabeads™ Protein G (10007D, Thermo Fisher Scientific) were washed two times with 0.5 mL of RIP buffer and then added to each sample, incubated for another two hours at 4 °C. The beads were collected and washed three times with 0.5 mL of RIP buffer; the last wash was performed with 0.5 mL of 1X PBS. Next, beads were resuspended in 0.1 mL of 1X PBS, followed by DNA digestion with 2U of DNase I (1 U/µL, EN0525, Thermo Fisher Scientific) at 37 °C for 15 min. Then, 0.1% SDS (Sigma) and 50 μg proteinase K (20 mg/mL, AM2548, Thermo Fisher Scientific) were added to each sample and incubated for 30 min at 50 °C (with shaking). DNA- and protein-free RNAs were collected from the beads using a magnetic stand. RNAs were further purified through RNA Clean and Concentrator-5 (R1013, Zymo Research) and eluted in 8 μL. Purified RNAs were used to make the RNA-seq libraries using NEBNext Ultra Directional RNA Library Prep Kit for Illumina (E7760, NEB), followed by RNA-seq. All the libraries were checked through a Bioanalyzer for quality control. Paired-End (PE) 150 bp reads were generated using a HiSeq-4000 sequencer (Illumina). RIP-seq was performed in duplicates.

### m^6^A-sequencing (m^6^A-seq or m^6^A RIP-seq)

m^6^A-seq or m^6^A RIP-seq was performed based on previously reported protocols^6, 9, 50^ with several modifications. Briefly, 5 μg of total RNA was chemically fragmented in 2 μL of 10X fragmentation buffer (100 mM Tris-HCl, 100 mM ZnCl_2_) in a total volume of 20 μL at 94 °C for 5 min. 2 µL of 0.5 M EDTA was immediately added to the reaction to stop the fragmentation, followed by purification and elution of fragmented RNA using oligo Clean and Concentrator Kit (D4061, Zymo Research). Purified fragmented RNA (usually ∼ 100 nt) was diluted in m^6^A–Binding Buffer (0.05 M Tris-HCl (pH 7.5), 0.150 M NaCl, 1% NP-40, 0.05% EDTA) and denatured for 5 min at 75 °C. After denaturing, fragmented RNA samples were mixed with 5 μg anti-m^6^A antibody (Cat. No. 202003, Synaptic Systems) and 40 U of RNase inhibitor (Thermo Fisher Scientific) and incubated for 1 hour at room temperature with gentle rotation. Next, 50 μL of Dynabeads™ Protein A/G (Thermo Fisher Scientific) were washed two times with 0.5 mL of m^6^A–Binding Buffer and then added to the ‘RNA-antibody mixture,’ incubated for two hours at 4 °C with rotation. ‘RNA-antibody-Dynabeads’ were pelleted using magnetic stand, washed for 3 min each with gentle rotation in the following order: twice with 500 µL of m^6^a-Binding Buffer (0.05 M Tris-HCl (pH 7.5), 0.150 M NaCl, 1% NP-40, 0.05% EDTA), once with low-salt buffer (0.25X SSPE, 0.001 M EDTA, 0.05% Tween-20, 37.5 mM NaCl), once with high-salt buffer (0.25X SSPE, 0.001 M EDTA, 0.05% Tween-20, 137.5 mM NaCl), twice with TET buffer (1X TE, 0.05% Tween-20). Fragmented RNA was eluted from the beads using elution buffer (0.02 M DTT, 0.150 M NaCl, 0.05 M Tris-HCl, 0.001 M EDTA, 0.1% SDS) in 50 µL for four times at 42 °C for 5 min with shaking. Eluted RNA (total 200 µL) was purified and concentrated using RNA Clean and Concentrator-5 kit (R1015, Zymo Research). 10 ng of purified fragmented total RNA (before adding the antibody/ IgG) was collected and used as an input. Both ‘m^6^A RIP-ed RNA’ and ‘total RNA’ (input) were subjected to RNA-seq library preparation using SMARTer Stranded Total RNA-Seq Kit – Pico Input Mammalian (Cat. No. 635005, TAKARA). All the libraries were analysed through a Bioanalyzer for quality control, and PE 150 bp reads were generated using a HiSeq-4000 sequencer (Illumina). m^6^A-seq was performed in duplicates.

### RNA-seq

Total RNA (DNA free) was isolated through RNeasy Mini Kit (Cat. No. 74104, Qiagen), followed by ribosomal RNA (rRNA) depletion using NEBNext rRNA Depletion Kit (E6310, NEB). The ribosomal-depleted RNA was used to make the RNA-seq libraries using NEBNext Ultra Directional RNA Library Prep Kit for Illumina (E7760, NEB). These libraries were analysed through a Bioanalyzer for quality control. PE 150 bp reads were generated using a HiSeq-4000 sequencer (Illumina). RNA-seq was performed in duplicates.

### RNA-seq for RNA decay/stability

0.5 million cells (per well of a 6-well plate) were plated of wild-type and *Mov10 KO* mESCs. 48 hours later, the mESC medium was replaced with a fresh medium containing 5 μg/mL actinomycin D. Following actinomycin D treatment, cells were collected at different time points (0, 3 and 9 hours) and washed twice with 1X PBS. The total RNA was extracted using *Quick*-RNA Miniprep kit (R1054, Zymo Research), followed by poly (A) mRNA isolation using NEBNext Poly(A) mRNA Magnetic Isolation Module (E7490, NEB). Prior to RNA-seq library construction, ERCC RNA spike-in control (4456740, Thermo Fisher Scientific) was added to each sample (0.1 µL per sample). RNA-seq libraries were prepared using the NEBNext Ultra Directional RNA Library Prep Kit (E7760, NEB). Quality control was checked through a Bioanalyzer, followed by high-throughput sequencing using a HiSeq-4000 sequencer (Illumina) to obtain 150 PE reads. Duplicates were used for each time point (0, 3 and 9 hours) from wild-type and *Mov10 KO*.

### Direct RNA-seq for poly (A) length measurement

4 µg of total RNA from wild-type and *Mov10 KO* mESCs were used. Poly (A) containing mRNAs were enriched with Oligo(dT)25 Dynabead™ (61002, Thermo Fisher Scientific) and libraries were prepared using the Direct RNA Sequencing Kit (SQK-RNA002, Oxford Nanopore Technologies). All the libraries were multiplexed using the DeePlexiCon adapters^51^, loaded to Nanopore R9 Flow Cells (FLO-MIN106D, Oxford Nanopore Technologies) and run on a MinION sequencer. Duplicates from wild-type and *Mov10 KO* were used.

### Ribo-seq profiling

Wild-type and *Mov10 KO* mESCs were plated on a 15 cm dish. Once cells reached 80% confluency (∼30 million cells), fresh mESC media with cycloheximide (100 µg/mL) was added to the cells and incubated at 37 °C for 10 min to arrest the translation. Then, plates (with cells) were immediately transferred on ice and cells were washed twice with ice-cold 1X PBS containing 100 µg/mL cycloheximide. 800 µL lysis buffer (30 mM Tris-HCl (pH 7.4), 600 mM NaCl, 30 mM MgCl_2_, supplemented with Protease Inhibitor Cocktail (1:1000 dilution, P8340, Sigma), 80 U of RNase inhibitor (20 U/µL, AM2696, Thermo Fisher Scientific), 0.8 µL of ß-mercaptoethanol (21985023, Thermo Fisher Scientific) and 100 µg/mL cycloheximide) was added to the cells and scraped from the plate. The cell lysates were transferred to clean 1.5 mL tubes and incubated at 4 °C for 10 min with gentle rotation. Next, cell lysates were centrifuged at 10,000 RPM for 10 min at 4 °C, and the supernatant was collected. 100 µL of the total cleared lysates were collected for ‘input’ (control), 10 µL of 10% SDS was added and purified through RNA Clean & Concentrator-25 Kit (R1017, Zymo Research) (following protocol for > 200 nt RNA). ‘Total RNA’ was eluted in 25 µL of nuclease-free water, and RNA-seq libraries were prepared using NEBNext Ultra Directional RNA Library Prep Kit (E7760, NEB).

400 μL of the total cleared lysates were treated with 4 μL of DNase I (1 U/μL, EN0525, Thermo Fisher Scientific) and 8 μL of RNase I (100 U/μL, AM2295, Thermo Scientific) for 45 min at room temperature with gentle shaking to obtain ribosome protected fragments (RPFs) RNA. 480 U of RNase inhibitor (20U /µL, AM2696, Thermo Fisher Scientific) was added to the lysates to quench the reaction, and samples were immediately transferred on ice. Next, RPFs RNA bound 80s monosomes (200 µL of RPFs nuclease digestion, out of 400 µL were used; the remaining 200 µL were snap-frozen in liquid nitrogen and stored at –80 °C) were isolated through MicroSpin S-400 columns (GE Healthcare, Cat. No. 27-5140-01), and 20 µL of 10% SDS were added; final volume was 220 µL. Further RPFs RNA were purified through RNA Clean & Concentrator-25 Kit (R1017, Zymo Research) using a modified protocol optimised for small RNA fragments as follows: a) in Zymo protocol Step 1, 220 μL Binding Buffer (from Zymo Kit) was used; b) in Zymo protocol Step 2, 495 μL absolute ethanol was used; c) purification was continued according to the manufacturer’s instructions; d) RPFs RNA samples were eluted in 60 μL of nuclease-free water. End repair of RPFs RNA samples was conducted through dephosphorylation reaction—by adding 7.5 μL T4 polynucleotide kinase (PNK) 10X buffer (B0201S, NEB), 1.5 μL ATP (100 mM, R041, Thermo Fisher Scientific), 1.5 μL RNaseOUT (10777019, Thermo Fisher Scientific), 1.5 μL T4 PNK (10U /µL, M020S, NEB) and 3 μL RNase-free water to a final volume of 75 μL. This was incubated at 37 °C for 1.5 hours, followed by column purification using RNA Clean & Concentrator Kit-5 (R1015, Zymo Research). End-repaired RPFs RNA were eluted in 20 µL of nuclease-free water, mixed with 10 µL of 2X RNA loading dye (R0641, Thermo Fisher Scientific), denatured at 70 °C for 3 min and separated on a 15% polyacrylamide TBE-urea gel in 0.5X TBE buffer alongside with small RNA marker (Cat. no. R0007; Abnova) and customed 26-nt and 32-nt ssRNA oligos. The gel was run for 1 hour at 150 V and stained for 10 min with 1:10,000 SYBR Gold in 0.5X TBE. The gel region between the 26–32 nt was excised with a clean, sterile scalpel; transferred to 1.5 tubes; crushed into small pieces and eluted in 300 μL of 0.3 M NaCl solution with 1 μL of RNaseOUT overnight at 4 °C on rotation. The gel slurry was transferred to 0.22 μm cellulose acetate membrane containing centrifuge tubes (Spin-X centrifuge tube filters, CLS8161-100EA, Corning) and centrifuged at a maximum speed for 5 min at 4 °C. The collected RNA filtrate was precipitated by adding 30 µL of 3 M sodium acetate, 1 µL glycogen, 800 µl ethanol through centrifugation at a maximum speed for 30 min at 4 ^°^C. The RNA pellet was washed with 80% ethanol once at a full speed for 10 min at 4 ^°^C. Finally, the RNA pellet was dried and dissolved in 12 µL nuclease-free water. ‘RPFs RNA’ libraries were prepared using a NEBNEXT Small RNA Library Prep Set for Illumina (E7330, NEB), following the manufacturer’s instructions. This experiment was performed in triplicates from wild-type and *Mov10 KO*.

### Immunoprecipitation of MOV10 for identification of its interacting proteins

Wild-type J1 mESCs were cultured in eight 15 cm dishes, and cells were harvested at 80–90% confluency. After two washes with 1x PBS, cells were lysed in 3 mL of NP-40 lysis buffer (150 mM of NaCl, 1% NP-40, 50 mM of Tris-HCl (pH 8), with freshly added protease inhibitor (1:200 dilution)) (3 volumes of the cell pellet size—cell pellet size was 1 mL and 3 mL of NP-40 lysis buffer were used) for 30 min on ice. The total cell lysate was cleared through centrifugation at 14,000 g for 10 min at 4 ^°^C, the supernatant was collected, and protein concentration was measured using Quick Start Bradford Protein Assay (5000201, Bio-Rad). Cleared cell lysate containing 5 mg of total protein was used for each IP. Either 10 µg of MOV10 antibody or IgG (control) were incubated with cell lysate (5 mg of total protein) for two hours at 4 °C with gentle rotation. 100 μL of Dynabeads™ Protein G (10007D, Thermo Fisher Scientific) were washed two times with 1 mL of NP-40 buffer and then added to ‘lysate-MOV10 antibody’ and ‘lysate-IgG’ mixture, incubated for another two hours at 4 °C with gentle rotation. Then beads were collected through a magnetic stand and washed three times with 1 mL of wash buffer (150 mM of NaCl, 50 mM of Tris-HCl (pH 8)), followed by elution three times (150 µL/elution) in 0.2 M glycine pH 2.5. Further, eluted samples were neutralised with a one-quarter volume of 1M Tris-HCl (pH 8) prior to in-solution digestion for mass spectrometry (MS). IP-MS of MOV10 was performed in duplicates.

For Western blot analysis, 1% of total cell lysate was collected as an ‘input’ prior to IP. MOV10/IgG IP-ed materials were collected at the last washing step of beads. 1 mL of wash buffer was added to the beads, resuspended and 100 µL of wash buffer with beads were collected through a magnetic stand. MOV10/IgG IP-ed samples were eluted from the beads in 30 µL of XT buffer (Cat. No. 1610791, Bio-Rad) through heating at 95 ^°^C for 10 min. Input and MOV10/IgG IP-ed samples were resolved on a 4–12% Criterion XT Bis-Tris Protein Gel (Cat. No. 3450125, Bio-Rad) and analysed through Western blot using specific antibodies.

### Protein co-immunoprecipitation of MOV10 to validate its interacting partners

For this purpose, we used MOV10-FB tagged mESC lines that expresses comparable levels as endogenous MOV10 protein, as mentioned earlier. MOV10-FB and BirA (control) mESCs were grown in a 15 cm dish and harvested once they reached 80–90% confluency. Briefly, cell pellets were washed twice in 1X PBS and dissolved in 1 mL of NP-40 buffer (150 mM of NaCl, 1% NP-40, 50 mM of Tris-HCl (pH 8), with freshly added protease inhibitor (1:200 dilution)) in the presence or absence of 200 U of RNase I (100 U/µL, AM2294, Thermo Fisher Scientific). 5% of the total cell lysate was collected as an ‘input’ before starting the IP. The remaining MOV10-FB and BirA (control) total lysate (950 µL) were incubated with 50 µL streptavidin Dynabeads™ (Cat. No. 65601, Thermo Fischer Scientific) for two hours at 4 °C. Subsequently, beads were collected through magnetic stand and washed three times with 0.5 mL of wash buffer (150 mM of NaCl, 50 mM of Tris-HCl (pH 8)) at 4 °C. IP-ed protein and its interacting partners were eluted from the beads in 60 µL of XT buffer (1610791, Bio-Rad) through heating at 95 °C for 10 min. Both input and IP-ed samples were resolved on a 4–12% Criterion XT Bis-Tris Protein Gel (Cat. No. 3450125, Bio-Rad) and analysed by Western blot using specific antibodies.

### LC-MS/MS of immunoprecipitated proteins

Immunoprecipitated samples were denatured and alkylated by adding TCEP (Tris(2-carboxyethyl) phosphine hydrochloride) and CAA (2-Chloroacetamide) to a final concentration of 10 mM and 40 mM, respectively, and the mixture was incubated at 55°C for 15 min. Sequencing grade trypsin was added at an enzyme to protein ratio of 1:50 and incubated overnight at 37°C after the proteins were trapped using S-Trap mini columns (Profiti). Tryptic peptides were sequentially eluted from the columns using (i) 50 mM TEAB, (ii) 0.2% formic acid and (iii) 50% acetonitrile, 0.2% formic acid. The fractions were pooled and concentrated in a vacuum concentrator prior to MS analysis.

Using a Dionex UltiMate 3000 RSLCnano system equipped with a Dionex UltiMate 3000 RS autosampler, the samples were loaded via an Acclaim PepMap 100 trap column (100 µm x 2 cm, nanoViper, C18, 5 µm, 100å; Thermo Scientific) onto an Acclaim PepMap RSLC analytical column (75 µm x 50 cm, nanoViper, C18, 2 µm, 100å; Thermo Scientific). The tryptic peptides were separated by increasing concentrations of 80% ACN / 0.1% FA at a flow of 250 nl/min for 128 min and analyzed either with an Orbitrap Fusion Tribrid or QExactive Plus mass spectrometer (Thermo Fisher Scientific) operated in data-dependent acquisition (DDA) mode. Acquired .raw files were searched against the murine UniProtKB/SwissProt database (v2017_07) using Byonic (Protein Metrics) considering a false discovery rate (FDR) of 1%. Carbamidomethylation of cysteine residues was specified as a fixed modification. Oxidation of methionine and acetylation of protein N-termini were selected as variable modifications. Trypsin was used as the enzymatic protease and up to two missed cleavages were allowed.

### Western Blot

Total cell lysate or IP-ed material were mixed with XT buffer (1610791, Bio-Rad), heated at 95 °C for 5 min and resolved on a 4–12% Criterion XT Bis-Tris Protein Gel (Cat. No. 3450125, Bio-Rad). Proteins on the gel were transferred to the PVDF membrane, and specific antibodies were used to detect proteins of interest.

### RIP-seq analysis

Adapters and low-quality sequencing reads < 30 bp of RIP-seq samples were filtered out using TrimGalore (v0.6.6). Clean reads of RIP and matched IgG samples were aligned to the mm10 mouse genome assembly and the GENCODE version M23 mouse transcriptome using STAR RNA-seq alignment program (v2.6.3b)^52^. RIP-seq enriched regions of each RIP sample were identified using the MACS2 peak-calling algorithm (v2.1.1.20160309)^53^ to their corresponding IgG sample as control. Fold enrichment was calculated as FC (RIP/IgG) > 2; *q*-values < 0.1.

### m^6^A-seq/ m^6^A RIP-seq analysis

TrimGalore (v0.6.6) was used to eliminate adapters and reads < 30 bp of m^6^A RIP-seq and input (RNA-seq), followed by alignment to the mm10 mouse genome assembly and the GENCODE version M23 transcriptome using STAR RNA-seq alignment program (v2.6.3b)^52^. Enriched m^6^A peaks were identified compared to its corresponding input control sample (i.e., RNA-seq), using ExomePeak R/Bioconductor Package^54^. m^6^A/A enrichment was calculated as fold change (m^6^A RIP/Input) > 1.5; *p*-values < 0.05. Coverage counts BigWig tracks were extracted from m^6^A RIP and input control samples using the ‘bamCoverage’ program from the deepTools software suite (v.2.5.3)^55^. The signal was smoothened using a sliding window approach.

Overlapping RIP-seq and m^6^A/A peaks were extracted using ‘bedtools intersect’ from the BEDTools software suite (v2.26.0)^56^ for further downstream analysis.

### RNA-seq analysis

Raw fastq reads were filtered through TrimGalore (v0.6.6) to remove adaptors and < 30 bp reads. Filtered reads were aligned to the mm10 mouse genome assembly and the GENCODE version M23 mouse transcriptome using the STAR RNA-seq alignment program (v2.6.3b). Read counts per transcripts were obtained from the aligned reads using htseq-count, part of the HTSeq package. Differential gene expression analysis was conducted using the DESeq2 (v1.14.1)^57^. Significantly altered transcripts (Log2-fold-change >< 0.5; Benjamini-Hochberg *p*-values < 0.05) were used for downstream analysis. MA and violin plots were created employing the ggplot2 R package (v3.4) that used log2 fold changes and mean normalised counts calculated by the DESeq2 R package. BigWig tracks were generated using the deepTools software suite (v.2.5.3) ^55^.

### Ribo-seq analysis

TrimGalore (v0.6.6) was used to discard both adapters and reads < 20 bp. Filtered data were uploaded directly to RiboToolkit workflow^58^. Also, the RNA-seq normalised read count matrix file (input) was uploaded to RiboToolkit for calculating TE. Briefly, RiboToolkit was used to distinguish RPFs (aligned to mRNA) from tRNA, rRNA and snRNA. RPFs mRNA (between 26–32nt) were used to calculate coding frames distribution, Ribo-seq quality evaluation, codon usage, reproducibility between the replicates and mRNA TE. TE was calculated using the ratio between RPFs and mRNA reads (from the RNA-seq, as an input) for each transcript. Differential TE between wild-type and *Mov10 KO* was calculated as log2 fold change, and ggplot2 package was used for cumulative plots.

### LC-MS/MS analysis

Total number of spectra of each identified protein was used for the analysis and log2 fold change (FC) was calculated between IP-ed sample and control. Significant enriched/interacting proteins were selected based on log2 FC (IP-ed sample/control) >1, *p-*values <0.05; others were considered as non-significant. Volcano and scattered plots were generated in R.

### Motif analysis

Peak annotation and RNA motif search were accomplished using the HOMER software (v4.11)^59^. Motifs were found within ± 200 nt of the peak centre of m^6^A.

### Statistical analysis

All statistical analyses for RIP-seq, m^6^A-seq/m^6^A RIP-seq, RNA-seq and Ribo-seq data were performed using the R statistical program (v3.3.2). Various R packages and GraphPadPrism9.0 were used to perform data analysis, such as one and two-way ANOVA test, t-test (one sample/ Wilcoxon test and paired t-test), PCA, correlation, clustering, heatmaps, etc. All the experiments were performed either in duplicates or triplicates and mentioned in the figure legends and/or method section. *p*-values < 0.05 were considered statistically significant; all *p*-values are indicated in the figure legends.

### mRNA decay (HL calculation)

150 paired-end (PE) RNA-seq reads in fastq format were aligned to mm10 genome using TopHat. Only concordant read pairs where each read had a mopping quality ≥ 50 are kept. Duplicates were removed with MarkDuplicates from Picard Tools. Aligned reads were assigned to genes usenf featureCounts from Rsubread R (Bioconductor) package. Log fold changes between 0 and 3 or 9 hours of wild-type and *Mov10 KO* were computed using edgeR R (Bioconductor) package.

Let *A(t)* be the expression of a mRNA at time *t* ≥ 0. Assuming the usual exponential decay model we obtain:

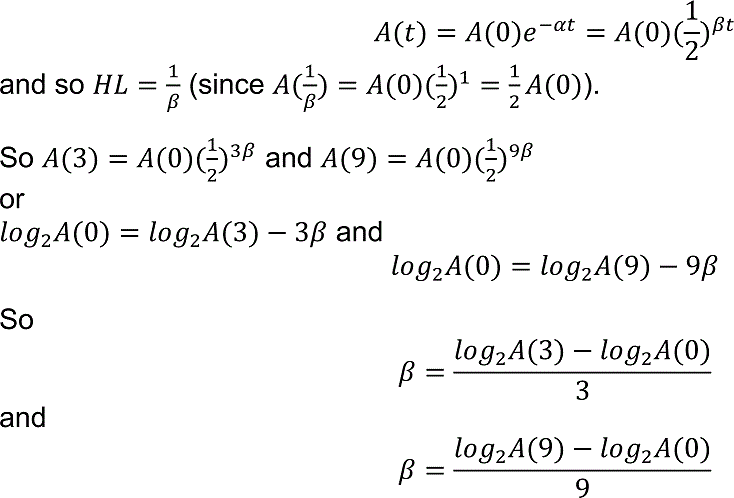

Previous reports computed:

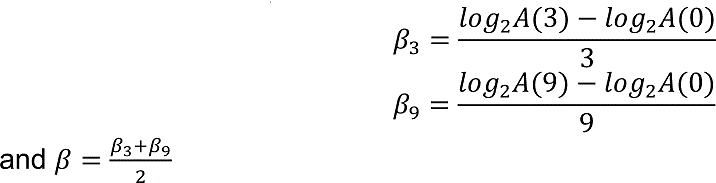

In reality, *log_2_A(0), log_2_A(3) and log_2_A(9)* are not measured exactly. It is quite common to assume that what we measure is:

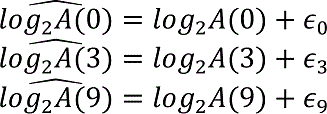

where ∈_0_, ∈_3_ and ∈_9_ are independent (normal) random variables with mean 0 and common variance σ^2^.

So in fact we have:

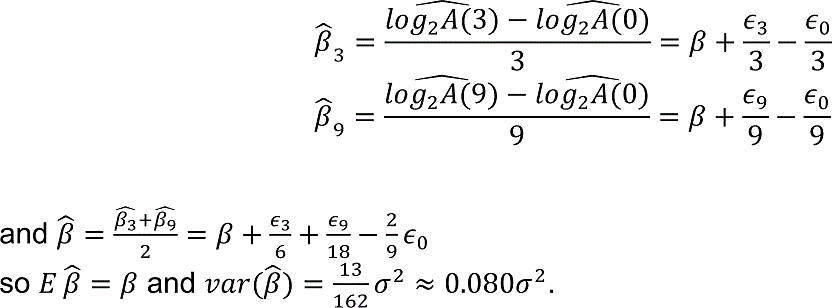

To obtain a better (in terms of variance) estimator we placed:

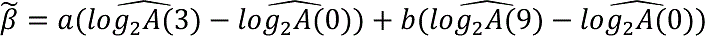

(1) where *a* and *b* are constants.

In order for β^∽^ to be an unbiased estimator of β we need:

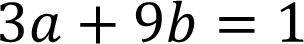

(2) It is also not hard to see that –

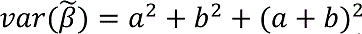

We are looking for *a* and *b* which minimise (2) subject to (1).

It follows from (1) that 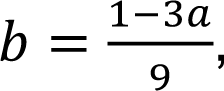 so (2) becomes

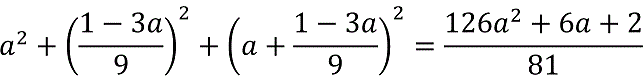

which is minimised for 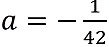 and then 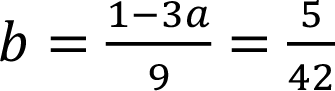

Using (2) we observed that in this case 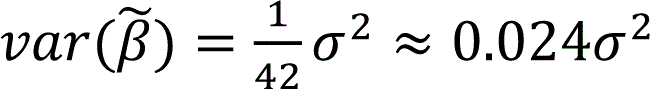, which is much smaller than that of 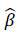.

## Direct RNA-seq bioinformatics analysis

Samples were demultiplexed with the DeePlexiCon pipeline^51^ using an accuracy threshold of > 0.95. We did the base calling with Guppy for each read, followed by alignment to the reference transcriptome with Minimap2^60^. Then, we indexed the based-called fastq files and mapped bam files with Nanopolish^61^ and SAMtools^62^, respectively. The poly (A) tail length was determined with Nanopolish.

## Splicing and m^6^A analysis

For splicing analysis, a splice junction BED file was generated using RegTools (v0.5.1)^63^ with a minimum intron size of 20 and a maximum intron size of 500,000 and a minimum anchor length of 6 nt. The BAM and BED files were taken as input for SpliSER (v0.1.2)^36^, along with mm10 gene annotation.

We plotted the cumulative frequency of high confidence splice sites with SSE ranging (0.1) that either did or did not overlap with m^6^A sites (+/- 300bp from GGm^6^ACU motif). High-confidence splice sites were defined as those with ≥ 10 reads overlapping the splice site in all samples and utilised at least 5% of the time (SSE > 0.05) in both replicates of at least one genotype. We tested the relationship between SSE and m^6^A sites overlap using a Chi-squared test, taking low- and high-SSE groups of sites (SSE < 0.5 and SSE > 0.5, respectively).

## Microscopy analysis

Imaging was performed using a Zeiss Airy Scan in 4Y mode using a 63x 1.4 NA objective with a stack spacing of 400 nm. Analysis was performed by particle analyser in ImageJ, and number of nucleus were counted manually. For co-localization analysis between MOV10 and EDC4, the locations of all pixels with MOV10 punctate structures were recorded and used as a primary. EDC4 channel was used as a secondary and were queried for presence of particles in the same locations recorded for MOV10.

## Data availability

All the high-throughput sequencing data from this study have been deposited in GEO under the accession numbers of GSE181821 and GSE181746.

## Acknowledgements

We thank the Monash Biomedical Imaging facility, Monash Mass Spectrometry facility and Genewiz high-throughput sequencing facility. This work is supported by the National Health and Medical Research Council (NHMRC) of Australia (APP1159461 and APP1182804) to P.P.D. S.K.B. is supported by Australian Research Council (ARC) DP19010479. M.M. is supported by the Intramural Research Program of the National Institutes of Health (NIH), National Institutes of Environmental Health Sciences (NIEHS) (ES103339-03). E.A.M is supported through Cancer Research UK (C13474/A18583, C6946/A14492) and the Wellcome Trust (104640/Z/14/Z, 092096/Z/10/Z).

## Author contributions

P.P.D. and P.T. conceptualised the study. M.M., D.B., D.H., Z.Z., H.G., A.F., H.C., S.A., R.S., S.C. and M.M. performed the experiments and analysed the data. Y.K., M.O., C.D. B.N.P. and M.M. performed bioinformatics analyses. M.M., Y.K., M.O., S.A., R.S., M.M., S.K.B., E.A.M., P.T. and P.P.D. interpreted the data. M.M., P.T. and P.P.D. wrote the manuscript.

## Competing interests

E.A.M is a co-founder and director of Strom Therapeutics, Cambridge, UK.

## Notes

### Competing Interest Statement

Eric miska is a co-founder and director of Strom Therapeutics, Cambridge, UK.

### Summary of Updates

Revised author list and affiliation

