## Supplemental information for "MOV10 facilitates messenger RNA decay in an *N6*-methyladenosine (m^6^A) dependent manner to maintain the mouse embryonic stem cells state"

### Extended Data Fig. 1

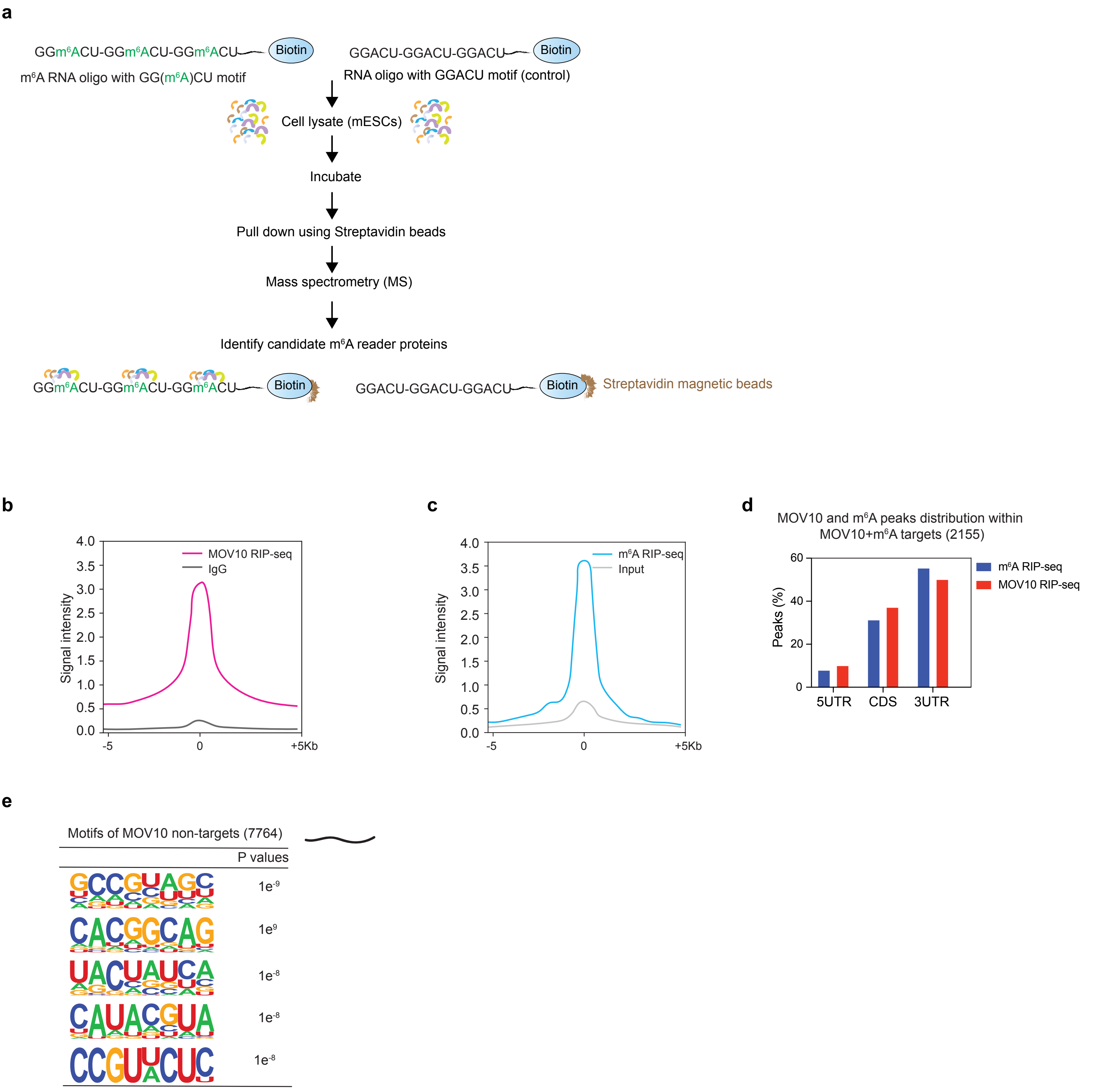

### Extended Data Fig. 2

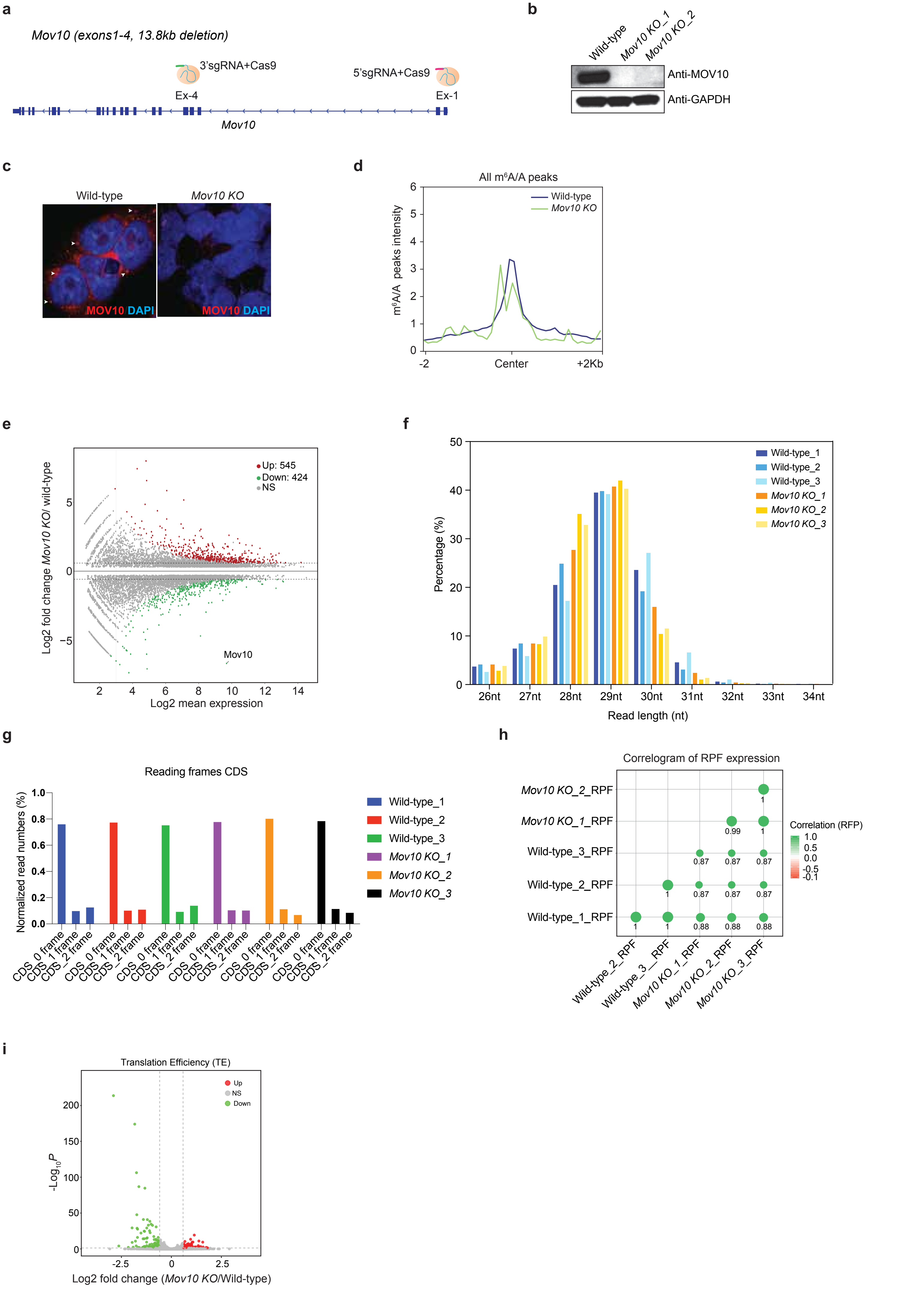

**Extended Data Fig. 3**

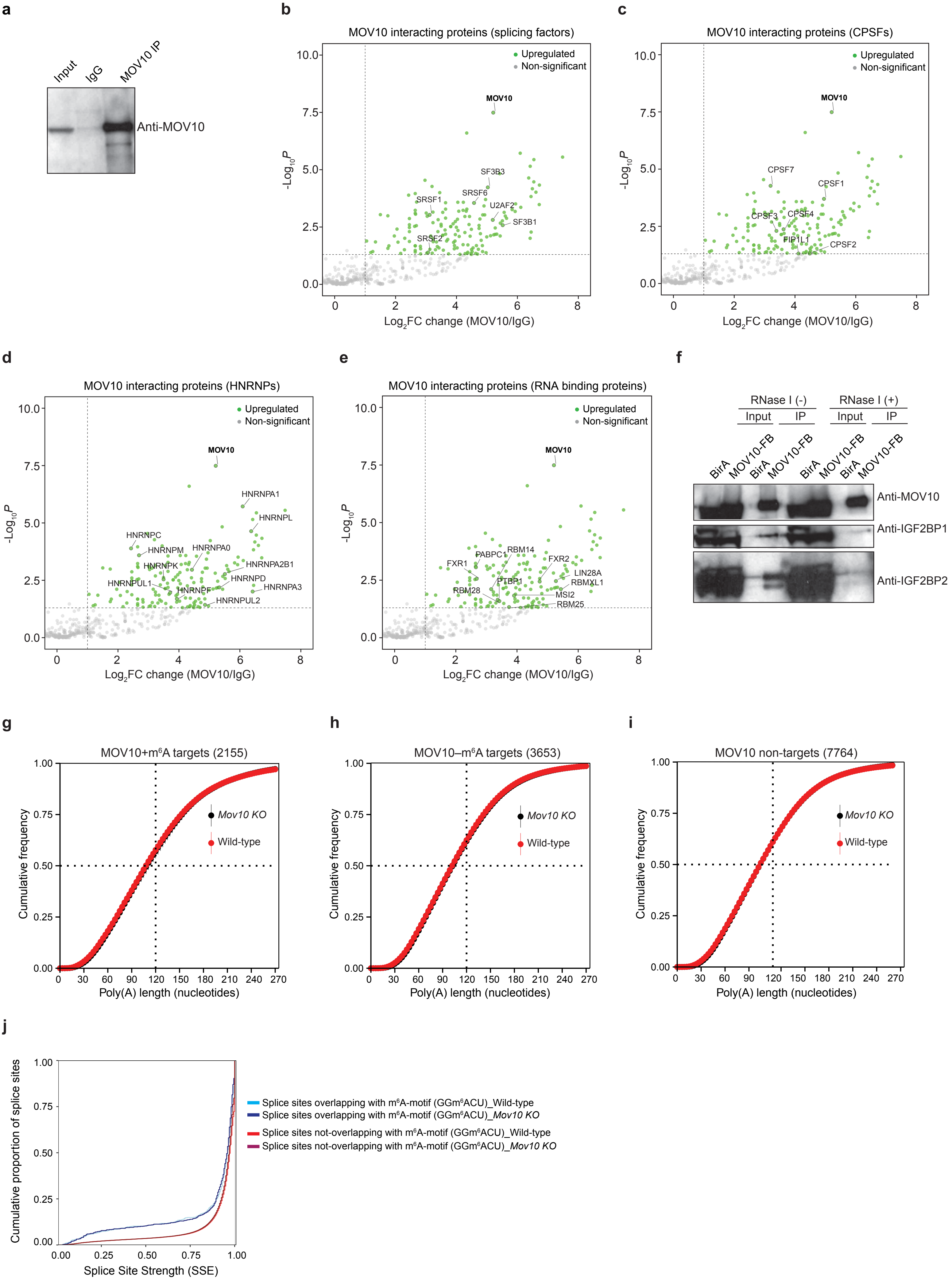

### Extended Data Fig. 4

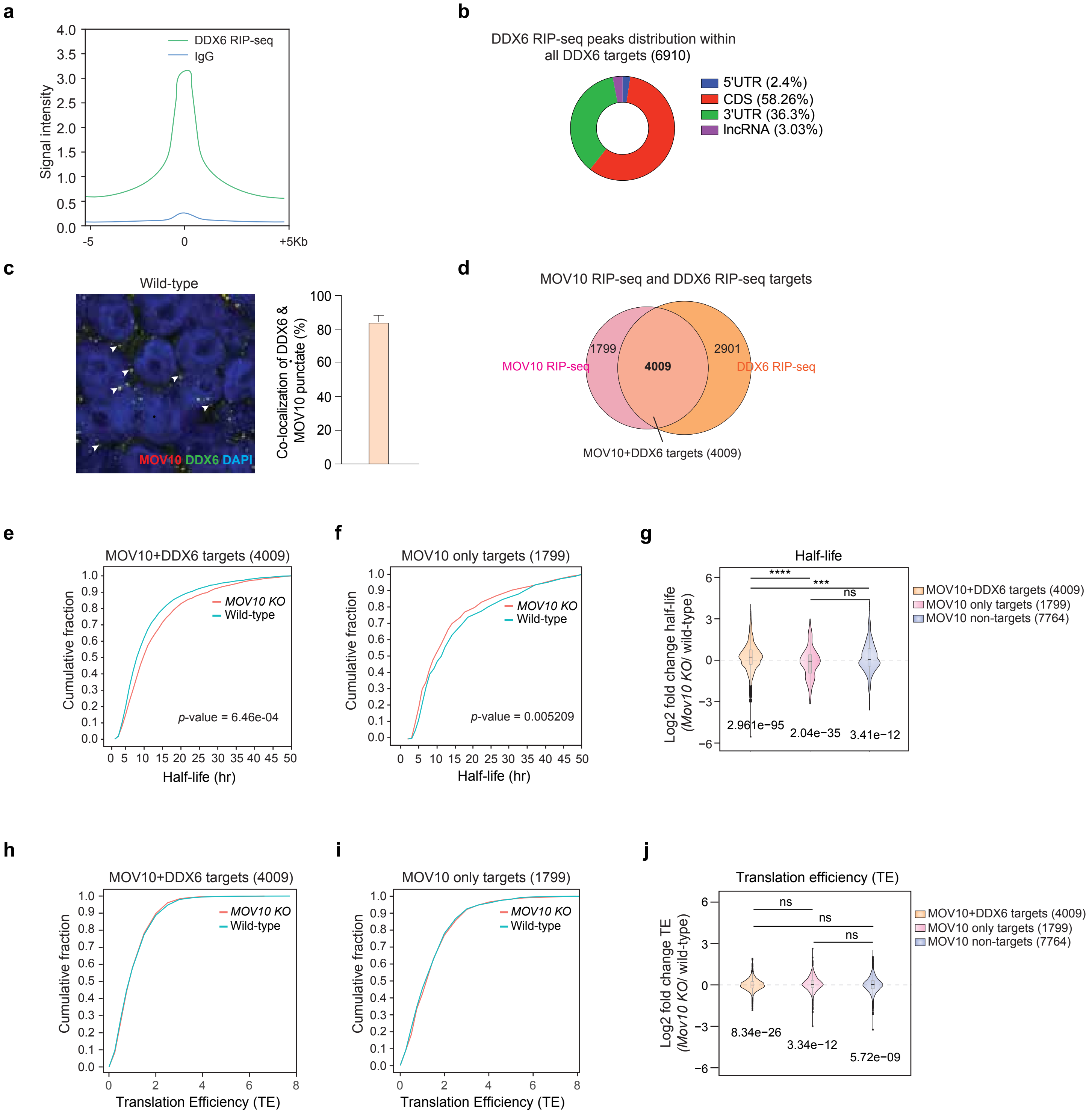

**Extended Data Fig.5**

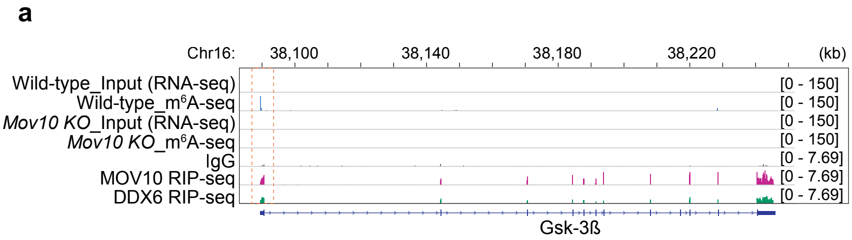


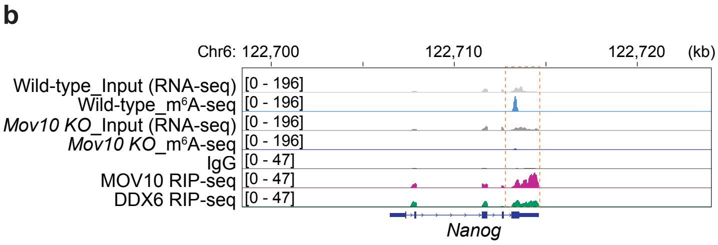

### Extended Figure Legends

**Extended Fig. 1. MOV10 identified as an m$^6$A-binding/reader protein that selectively binds to m$^6$A-containing mRNAs.**

a) A Schematic diagram illustrate the strategy to identify potential new m$^6$A-reader proteins in mESCs. Biotinylated single-stranded (ss) RNA probes – either containing 3 repeats of an m$^6$A consensus motif of GGm$^6$ACU (with methylated adenosine) or 3 repeats of GGACU (with unmethylated adenosine, used as a control) – were incubated with mouse embryonic stem cells lysate and streptavidin beads were used to pull down the RNA probes-bound proteins, which were subsequently identified through quantitative mass spectrometry-based proteomics. The possible candidate m$^6$A-reader proteins were identified based on their enriched binding to the methylated GGm$^6$ACU RNA probe over the unmethylated GGACU RNA probe. b) Profile plot showing RIP-seq signal intensities of MOV10 and IgG (control). c) Profile plot displaying RIP-seq signal intensity of m$^6$A, RNA-seq used as control/input. d) Percentage of MOV10 and m$^6$A peaks distribution within MOV10+m$^6$A targets. e) Top five motifs are shown of MOV10 non-targets (7,764) through HOMER analysis.

**Extended Fig. 2. MOV10 destabilises its mRNA targets in an m$^6$A-dependent manner.**

a) Schematic diagram demonstrating the strategy to generate *Mov10* homozygous/biallelic deletion or *KO* clones. Paired sgRNAs (5' and 3' sgRNAs) flanking exons 1-4 of *Mov10* were introduced to create homozygous deletion. b) Western blot result confirms successful generation of *Mov10 KO* mESC lines. c) Immunostaining of MOV10 display its sub-cellular localisation in wild-type that disappear in *Mov10 KO* mESCs, confirming *Mov10 KO* as well. d) m$^6$A/A peak intensities of 'all' m$^6$A-containing targets in *Mov10 KO* and wild-type. e) MA plot show overall gene expression changes in *MOV10 KO* compared to wild-type, obtain from RNA-seq. f) Ribosome profiling (Ribo-seq) display percentage (%) of ribosome protected fragments (RPFs) length distribution in a range of 26-34nt mRNA from wild-type and *Mov10 KO*. The majority of the RPFs peaks are shown in 28-29nt range (as expected), reflecting the size of a translating ribosome on the mRNAs. g) RPFs distribution on different coding frames (0, 1, 3) of CDS, showing majority of the RPFs are in frame 0 from wild-type and *Mov10 KO*. h) Ribo-seq correlogram demonstrate RPFs expression among wild-type and *Mov10 KO*, including their replicates. i) A volcano plot represents overall translation efficiency (TE) changes in *Mov10 KO* compared to wild-type.

**Extended Fig. 3. MOV10 interactome ascertains diverse protein networks of poly (A) length maintenance and splicing of mRNAs; however, MOV10 does not participate in regulating poly (A) length and splicing of mRNAs.**

a) Western blot showing MOV10 enrichment in MOV10 immunoprecipitation (IP). b-e) Different categories of significantly enriched MOV10-interacting proteins are presented (in green) (Log2FC>1; P-values <0.05). Non-significant proteins are labelled in grey. n=2 replicates. f) Co-immunoprecipitation of using streptavidin beads against biotinylated MOV10-FB tagged protein from mESCs extract in the presence or absence of RNase I, followed by Western blot analysis to detect MOV10 interacting partners. g-i) Cumulative frequencies of poly (A) length of MOV10+m$^6$A targets (2,155), MOV10-m$^6$A targets (3,653) and MOV10 non-targets (7,764) in *Mov10 KO* and wild-type. j) Splice sites strength estimates (SSE) are calculated around the +/- 300bp of m$^6$A sites (resides in GGm$^6$ACU motif), which show significant correlation between individual m$^6$A sites and individual splice-site strengths. However, no alternations of splice-site strength/usage were observed around the $m^6A$ sites in *MOV10 KO* compared to wild-type.

**Extended Fig. 4. MOV10 destabilises its $m^6A$-containing mRNA targets mainly within the P-bodies.**

a) Profile plot showing RIP-seq signal intensities of DDX6 and IgG (control). b) Pie-chart distribution of DDX6 binding sites at 5'UTRs, CDS and 3'UTRs and long non-coding RNAs (lncRNAs) within all DDX6 targets (6,910), obtained from DDX6 RIP-seq. c) Immunostaining showing sub-cellular co-localisation of MOV10 and DDX6 (as a P-body marker) as punctate structures. A bar chart represent quantification of co-localised of MOV10 and DDX6 punctates. d) A Venn diagram represent overlap targets (4,009) between – total MOV10 (5,808) and DDX6 (6,910) targets. e, f) Cumulative plots presenting HLs of MOV10+DDX6 targets (4,009) (e) and MOV10 only targets (1,799) (f) in wild-type v. *Mov10 KO*. Only significant *p*-values that represent the HL changes in *Mov10 KO* v. wild-type are shown in each graph. n=2 replicates. g) A violin plot exhibiting log2 fold change of HLs of MOV10+DDX6 targets (4,009), MOV10-only targets (1,799) and MOV10 non-targets (7,764) in *Mov10 KO* compared to wild-type. *P*-values are shown of each set of targets using Wilcoxon t-test (at the bottom). Significant alterations among two sets of targets represented through *p*-values (ANOVA test) (at the top); $p^{****}<0.0001$, $p^{***}<0.001$, ns= non-significant; n=2 replicates. h, i,) Cumulative plots demonstrating TE of MOV10+DDX6 targets (4,009) (h), MOV10 only targets (1,799) (i) in wild-type v. *Mov10 KO*. j) A violin plot showing log2 fold change of TE of MOV10+DDX6 targets (4,009), MOV10 only targets (1,799) and MOV10 non-targets (7,764) in *Mov10 KO* compared to wild-type.

*P*-values are indicated of each set of targets (at the bottom). Alterations among two sets of targets are ns= non-significant (ANOVA test); n=3 replicates.

**Extended Fig. 5. MOV10 destabilise Gsk-3ß mRNA in an $m^6A$-dependent manner to control the WNT/ß-CATENIN-NANOG regulatory pathway for the mESC state maintenance.**

a, b) $m^6A$ RIP-seq and RNA-seq (control) tracks showing the enrichment of $m^6A$ in mRNA targets (Gsk-3ß and Nanog). The RIP-seq of MOV10, DDX6 and IgG (control) tracks displaying enriched binding of MOV10 and DDX6 to their mRNA targets. Dotted box highlight overlapping $m^6A$ and MOV10, DDX6 enriched sites in mRNA targets, where $m^6A$ levels are reduced in *Mov10 KO* compared to wild-type.
